# The Aging Brain and Executive Functions Revisited: Implications from Meta-Analytic and Functional-Connectivity Evidence

**DOI:** 10.1101/2020.07.15.204941

**Authors:** Marisa K. Heckner, Edna C. Cieslik, Simon B. Eickhoff, Julia A. Camilleri, Felix Hoffstaedter, Robert Langner

## Abstract

Healthy aging is associated with changes in cognitive performance including executive functions (EFs) and their associated brain activation patterns. However, it has remained unclear which EF-related brain regions are affected consistently, because the results of pertinent neuroimaging studies and earlier meta-analyses vary considerably. We, therefore, conducted new rigorous meta-analyses of published age differences in EF-related brain activity. Out of a larger set of regions associated with EFs, only left inferior frontal junction (IFJ) and left anterior cuneus/precuneus (aC/PrC) were found to show consistent age differences. To further characterize these two age-sensitive regions, we performed seed-based resting-state functional connectivity (RS-FC) analyses using fMRI data from a large adult sample with a wide age range. We also assessed associations of the two regions’ whole-brain RS-FC patterns with age and EF performance. Although functional profiling and RS-FC analyses point towards a domain-general role of left IFJ in EFs, the pattern of individual study contributions to the meta-analytic results suggests process-specific modulations by age. Our analyses further indicate that left aC/PrC is recruited differently by older (compared to younger) adults during EF tasks, potentially reflecting inefficiencies in switching the attentional focus. Overall, our findings question earlier meta-analytic results and suggest a larger heterogeneity of age-related differences in brain activity associated with EFs. Hence, they encourage future research that pays greater attention to replicability, investigates age-related differences in deactivation, and focuses on more narrowly defined EF subprocesses, combining multiple behavioral assessments with multi-modal imaging.

**Highlights:** - Healthy aging is linked to deterioration in executive functions (EFs)

- ALE meta-analyses examined consistent age differences in brain activity linked to EFs

- In a larger set of EF regions, only left IFJ and (pre)cuneus were sensitive to age

- Advanced age was linked to weaker functional coupling within EF-related networks

- Our findings question earlier meta-analytic findings

## 1. Introduction

### 1.1 Executive Functions

Executive functions (EFs) are a loosely defined set of cognitive control processes that are taken to be critical for goal-directed thought and behavior in complex environments. Despite the lack of a clear formal definition of EFs as well as their ambiguous mapping on typical EF-tasks, there is relative agreement on their importance for regulating human behavior through modulating cognition in a top-down fashion (Diamond, 2013; Jurado & Rosselli, 2007). Different lines of research on how EFs might be best fractionated into subcomponents suggest models that argue for the existence of three core EFs: inhibitory control, working memory, and cognitive set shifting (e.g., Lehto, 1996; Miyake et al., 2000; for reviews see: Alvarez & Emory, 2006; Diamond, 2013). We acknowledge, however, that this differentiation is not undisputed (Baddeley & Hitch, 1974; Engle & Kane, 2004; Norman & Shallice, 1986; Stuss, 2006).

For a long time, it was thought that EFs were exclusively based on frontal lobe functioning as patients with frontal lesions often showed deficits in EFs leading to the interchangeable use of the terms “executive dysfunction” and “frontal lobe dysfunction” (e.g., Duncan, 1986; Owen et al., 1990; Shallice et al., 1982). However, patients with frontal lesions can perform within a normal range on EF tasks (e.g., Eslinger & Damasio, 1985; Shallice & Burgess, 1991) and patients with non-frontal lesions can show similar deficits like patients with frontal lesions (e.g., Anderson et al., 1991; Mountain & Snow, 1993). Years of research led Don Stuss and his collaborators (1995; 2006; 2011) to the assumption that there is a substantial fractionation of frontal lobe functions and that EFs represent only one functional category within the frontal lobes. Previous neuroimaging studies have revealed notable differences in the brain regions involved in EFs, which may be partly due to the elusive conceptualization of EFs (Collette et al., 2006) as well as the wealth of different perspectives, operationalizations, and traditions in this research, which have resulted in a co-existence of rather diverse labels for the brain networks and regions associated with EFs (Camilleri et al., 2018). Although there are differences between different tasks probing EFs, there also seem to be core regions consistently involved, like left inferior frontal junction (IFJ; e.g., Emery et al., 2008; Milham et al., 2002; Zysset et al., 2007). Duncan and collaborators (Duncan, 2010; Duncan & Owen, 2000; Fedorenko et al., 2013) investigated and defined these core regions and proposed that a “multiple-demand” (MD) brain system was consistently recruited during all kinds of cognitively demanding tasks.

Müller et al. (2015) used a similar approach: They integrated results from three neuroimaging meta-analyses investigating working memory (Rottschy et al., 2012), vigilant attention (Langner & Eickhoff, 2013), and inhibitory control (Cieslik et al., 2015), highly discussed subcomponents of EFs (Alvarez & Emory, 2006; Miyake et al., 2000), to define a common core network for EFs. This network was similar to Duncan’s MD system and comprised seven regions: mid-cingulate cortex/supplementary motor area (MCC/SMA), bilateral IFJ/inferior frontal gyrus (IFG), right middle frontal gyrus (MFG), bilateral anterior insula (aIns), right inferior parietal cortex (IPC), and intraparietal sulcus (IPS). Camilleri et al. (2018) went on to propose an extended MD network (eMDN) based on task-dependent and task-independent functional connectivity (FC) analyses seeded from the regions of the meta-analytically defined MD network by Müller and colleagues, to consider the perspective of a more widely distributed network. Camilleri et al. reported 17 regions as part of the eMDN (bilateral IFJ, aIns, SMA/pre-SMA, IPS, putamen, thalamus, MFG extending into the inferior frontal sulcus, dorsal premotor cortex [dPMC], and left inferior temporal gyrus). While the current paper focuses on EF-related activations, for the sake of completeness, we consider it necessary to briefly mention the functional relevance of the default-mode network (DMN) as it is assumed to be activated during stimulus-independent or spontaneous cognition and deactivated during externally focused cognition. The DMN comprises a network of brain regions that includes the precuneus (PrC)/posterior cingulate cortex (PCC), anterior medial prefrontal cortex (mPFC), and lateral inferior parietal cortex (Shulman et al., 1997; for reviews see: Anticevic et al., 2012; Raichle, 2015). In the interest of space, we did not go into more detail and referred to reviews on this topic.

Taken together, EFs seem to be a macro-construct rather than a single process, which involves distributed networks instead of any particular region, with a core network and more specific regions that are recruited depending on certain task demands (Camilleri et al., 2018; Miyake & Friedman, 2012; Teuber, 1972).

### 1.2 Healthy Aging

Healthy aging is associated with altered cognitive performance and brain activation patterns in several cognitive domains, especially non-routine tasks that tax executive control processes (Drag & Bieliauskas, 2010; Park et al., 2002; Stuss & Craik, 2019). Although the aging brain faces unfavorable changes, such as the decline of dopaminergic receptors (Li et al., 2001; Yang et al., 2003), volumetric shrinkage of many grey-matter structures (Raz et al., 2005; Resnick et al., 2003; Salat et al., 2004), and reduced white-matter density (Head et al., 2004; Wen & Sachdev, 2004), it also seems to aim for an allostatic maintenance of cognitive functions through functional reorganization. This indicates that the neurobiological substrates of our cognitive system are highly dynamic and adaptive across the lifespan (Greenwood, 2007; Park & Reuter-Lorenz, 2008).

A common finding is the reduced lateralization of brain activation in older adults, which is thought to be compensatory as it is correlated with better performance in older adults (“hemispheric asymmetry reduction in older adults” [HAROLD]; Cabeza, 2002). Furthermore, brain activation shifts from posterior to more anterior brain regions have been observed (“posterior– anterior shift in aging” [PASA]; Davis et al., 2007), which might be caused by older adults’ need for exerting executive control for previously automated operations. Additionally, it has been hypothesized that age-related cognitive and behavioral changes are associated with less specialized brain responses and a decrease in FC with age, in the context of structural and neurobiological changes as well as external experiences (Baltes & Lindenberger, 1997; Goh, 2011; Li & Sikström, 2002; Park et al., 2001; 2004). Reuter-Lorenz and Cappell (2008) postulated that the oft-reported increase in task-related lateral PFC activity with age compensates for less efficient neural circuits (“compensation-related utilization of neural circuits hypothesis” [CRUNCH]). Finally, Park and Reuter-Lorenz (2008) attempted to unite previous theories in their “scaffolding theory of aging and cognition” (STAC). In this context, “scaffolds” describe a supportive framework that helps maintain cognitive and behavioral performance at a relatively high level through advanced age via the strengthening of existing connections, development of new connections, and disuse of connections that have become fragile or deficient. These changes, in turn, are assumed to lead to increased bilateral and/or frontal activation in older adults.

Results from neuroimaging studies on age-related changes in the EF subcomponents working memory, inhibitory control, and cognitive flexibility are rather ambiguous. While some studies reported an increase in bilateral prefrontal activity (e.g., Emery et al., 2008; Madden et al., 1999) and a decrease in occipital activity (e.g., Ansado et al., 2012; Madden et al., 2002, 2010), other studies reported occipital activity increase (e.g., Bloemendaal et al., 2016; Van Impe et al., 2011) and frontal activity decrease in older adults (e.g., Bloemendaal et al., 2016; Schulte et al., 2011). Moreover, the age-related reduction in hemispheric asymmetry of brain activity is not found consistently (e.g., Carp et al., 2010; Toepper et al., 2014). This large amount of heterogeneous, partly contradictory findings illustrates the need for a quantitative data synthesis by means of meta-analysis in combination with taking a systems-level perspective, which includes identifying the connectional profiles of the identified regions with respect to the rest of the brain.

So far, three quantitative neuroimaging meta-analyses investigating cognitive aging in EFs have been published, each with their own limitations as discussed below: Spreng et al. (2010) performed an analysis across all then available experiments probing EFs in age, such as working memory, task switching, and inhibitory control. The authors found consistent EF-related increases in activity with age in bilateral dorsolateral prefrontal cortex (DLPFC), right posterior MFG/frontal eye field (FEF), left SMA, and left rostrolateral PFC as well as consistent decreases in activity with age in right ventrolateral PFC. Next, Turner and Spreng (2012) conducted separate meta-analyses for the EF subcomponents working memory and inhibition and found domain-specific patterns of across-experiment convergence. For working memory, consistent increases in activity with age were found in bilateral SMA, right MFG, left IFG, and left IPS; consistently lower activity in older adults was found in right IPS, left aIns/frontal operculum, and left FEF. For inhibition-related brain activity, consistent increases in activity with age was found in right MFG/IFG and left superior frontal gyrus, whereas consistent decreases in activity with age was found in right inferior occipital gyrus. Finally, a third meta-analysis by Di et al. (2014) found consistent increases in EF-related activation with age in bilateral IFG, left anterior cerebellum, left fusiform gyrus (FG), right MFG, and right parahippocampal gyrus. Consistently lower EF-related activation with age was found in bilateral Ins, left MFG, left medial frontal gyrus, and right MCC.

Taken together, the picture produced by these meta-analyses is largely inconclusive. This inconsistency across meta-analyses might result from methodological differences, such as the in- or exclusion of region-of-interest (ROI) contrasts, the particular selection of tasks included, or the approach to testing for age-related differences. Furthermore, all previous meta-analyses corrected for multiple comparisons by controlling the voxel-level false discovery rate (FDR), which has recently been shown to feature low sensitivity and a high susceptibility for false-positive findings in Activation Likelihood Estimation (ALE) meta-analysis (Eickhoff et al., 2016). In light of these inconsistencies and limitations of earlier efforts as well as the continued growth of the pertinent literature since 2014, a fresh meta-analysis on age-related differences in brain activity associated with EFs appeared much warranted.

### 1.3 Current Study

In a first step, coordinate-based ALE meta-analysis (Eickhoff et al., 2009, 2012; Turkeltaub et al., 2002, 2012) was used to synthesize results from neuroimaging studies investigating EFs in young and old participants. We started with a meta-analysis of within-group findings, pooling across experimental results obtained in young or old participants, respectively. This approach should test for consistent general EF-related brain activity in our sample of experiments, without regard to age-related differences. It was aimed at replicating previous findings of brain regions involved in EFs. Subsequently, we conducted further meta-analyses of published between-group contrasts, investigating consistent age differences in EF-related brain activity. As a methodological improvement over previous ALE meta-analyses on this topic, we used cluster-level family-wise error (FWE) correction for multiple comparisons (rather than FDR-based correction) and a minimum number of n = 17 experiments per analysis, following the recommendations by Eickhoff et al. (2016).

In a second step, task-independent whole-brain FC patterns of resulting age-sensitive regions were analyzed using resting-state (RS) functional magnetic resonance imaging (fMRI) data of healthy adults. Finally, we assessed the associations of the regions’ whole-brain FC with age and performance scores representing EF and its subcomponents in order to gain further insights into the mechanisms underlying cognitive aging.

In summary, this study aimed to investigate (i) which brain regions show consistent age differences in EF-related activity at the meta-analytic level, (ii) the connectional profiles of these age-sensitive regions, and (iii) how the connectivity profiles of these regions are affected by aging and EF-capacity.

## 2. Methods

### 2.1 ALE Meta-Analysis

#### 2.1.1. Sample

##### 2.1.1.1. Search for Studies

Pertinent studies were searched for in the databases Web of Science, PubMed (https://www.ncbi.nlm.nih.gov/pubmed/), PsycINFO (http://ovidsp.tx.ovid.com), and Google Scholar (http://scholar.google.de) using the following search strings: (1) title: “age” or “aging” or “ageing” or “age-related” or “older adults” or “old adults” or “life-span” or “elderly adults”; and (2) title: “executive functions” or “working memory” or “inhibition” or “cognitive flexibility”; and (3) abstract: “fMRI” or “functional magnetic resonance imaging” or “PET” or “positron emission tomography” or “neuroimaging” or “cerebral blood flow.” Subsequently, specific EF-related task labels were included in the search string as follows: for working memory, “n-back” or “Sternberg” or “delayed match* to sample” or “delayed simple matching”; for inhibitory control, “Stroop” or “flanker” or “Simon” or “stimulus-response compatibility” or “stop signal” or “go/no-go” or “stimulus detection” or “stimulus discrimination” or “selective attention”; and for cognitive flexibility, “task switching” or “dual task” or “set shifting.” The search criteria were partially motivated by previous meta-analyses regarding aging and EFs (Di et al., 2014; Spreng et al., 2010; Turner & Spreng, 2012). The decision on which tasks to include in the extended search string was made based on Diamond’s (2013) definition of typical tasks for each of the subcategories. Finally, earlier meta-analyses on this topic, reviews, and the reference lists of identified studies were inspected for additional studies to be included.

##### 2.1.1.2. Inclusion and Exclusion Criteria

We included only peer-reviewed publications of fMRI or positron emission tomography (PET) experiments performed in healthy young and old participants without any pharmacological manipulations or other extraneous interventions. Results of group analyses needed to be reported as coordinates of a standard reference space, that is, MNI (Montreal Neurological Institute) or Talairach (Talairach & Tournoux, 1988) space. Studies were only included if the whole brain was covered (i.e. coverage of at least 8 cm in the z-dimension). Consequently, no ROI-based results were included. However, some of the experiments we included reported masking of the between-group contrast with the task-positive main effect to restrict group differences to task-related regions (these studies are marked in Tables A1 and A2). We included results from contrasts between task and sensorimotor control or resting-baseline conditions, contrasts between different levels of task difficulty, as well as correlations between age and task-related activity.

Thus, deactivation data, results from connectivity analyses, or correlations and interactions with other variables (e.g., group × performance interactions, correlations with reaction time, etc.) were not considered. In case of uncertainty as to any of these criteria, the corresponding author of the given study was contacted for clarification (these studies are marked in Tables A1 and A2).

To minimize the risk that meta-analytic results were unduly biased by a particular publication, the contribution from any given study was limited to one experiment. If a study reported several experiments eligible for inclusion, their findings (i.e., reported coordinates) were pooled to constitute a single experiment, as suggested by Turkeltaub et al. (2012). Further, if contrasts for both transient and sustained brain activity were available, the contrast reflecting transient activity was chosen as it typically allows for a more process-specific interpretation. For the current approach, coordinates of within-group (i.e., main task effect per group) and between-group (i.e., contrast of task effects between groups: [young > old, old > young]) contrasts as well as correlations between task performance and age were included.

##### 2.1.1.3. Studies Included

After an initial screening of publication abstracts for topicality, 147 studies were retrieved in total. Applying the above criteria left us with 31 eligible studies reporting within-group task effects: 11 for working memory, 12 for inhibition, and 9 for cognitive flexibility. Of note, the study by Townsend et al. (2006) contributed results to two subdomains (inhibition and cognitive flexibility).

In the meta-analyses of age-related differences, 46 eligible studies were included in total: 15 for working memory, 19 for inhibition, 14 for cognitive flexibility, and 1 not clearly assignable to any subdomain. For clarification, not all studies included both within- and between-group contrasts, leading to somewhat different numbers of studies included in the within- and between-group meta-analyses, respectively. Of note, Eich et al. (2016) and Townsend et al. (2006) were included for inhibition and cognitive flexibility, and Lamar et al. (2004) for inhibition and working memory. Studies reporting different tasks (i.e., experiments that contribute to different subdomains) were pooled by the respective subdomain (vs. by subject group) and may thus contribute with two data points. To make sure that this pooling by subcomponent would not have an effect on the results, we additionally computed the meta-analyses with solely one point of data per study, i.e., pooling by subject group. The results were the same. For the sake of interpretability, we therefore decided to pool the aforementioned studies by subcomponent. See Figure 1 for an overview of the different analysis steps conducted.

**Figure 1.**
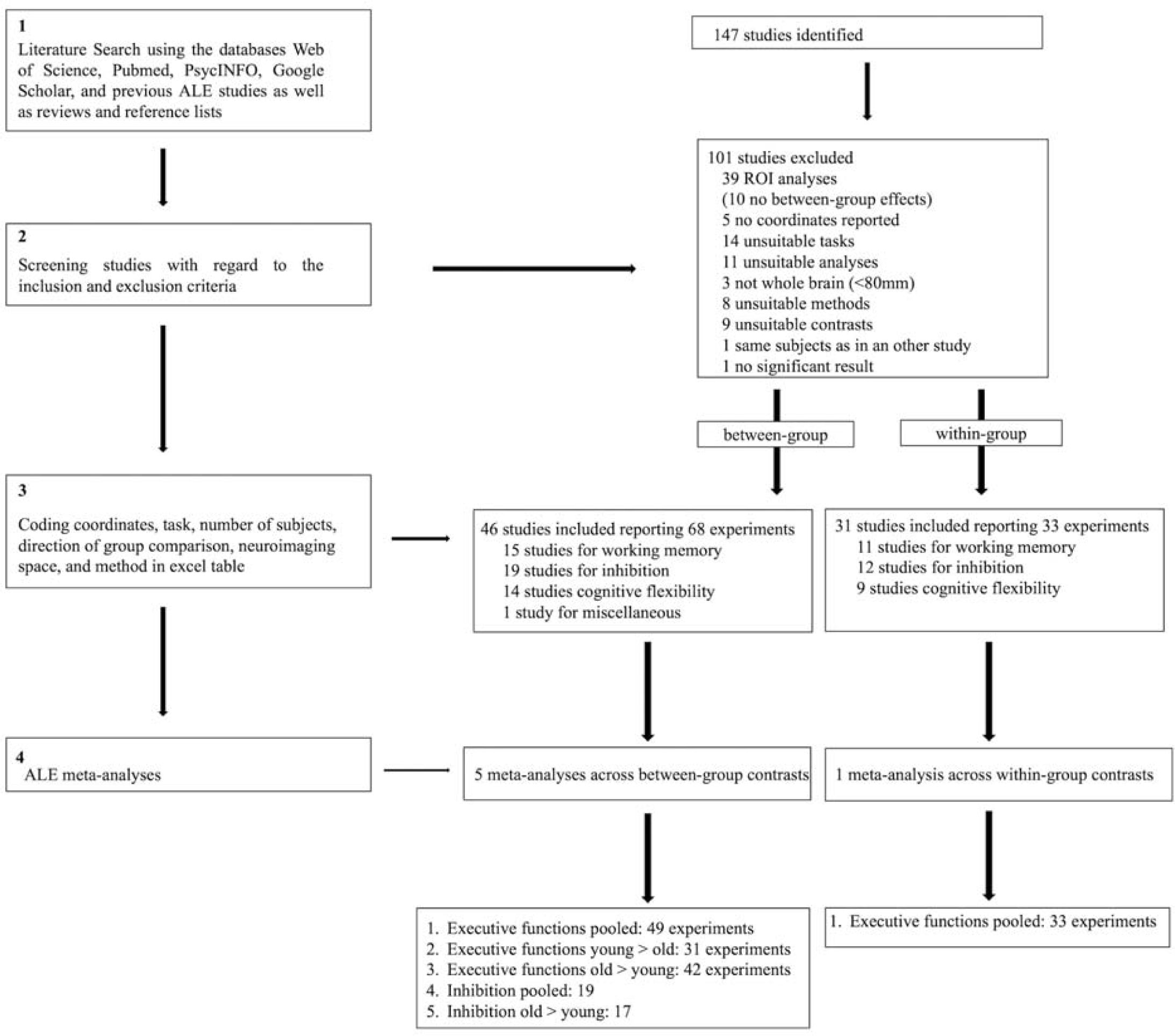
Flowchart of the meta-analysis steps conducted.

For further information about the studies included, please see Tables A1 and A2. A checklist for neuroimaging meta-analyses as recommended by Müller et al. (2018) can be found in Table A3.

#### 2.1.2. Activation Likelihood Estimation

##### 2.1.2.1. ALE Algorithm

All meta-analyses were conducted using the revised version of the ALE algorithm for coordinate-based meta-analysis of neuroimaging results (Eickhoff et al., 2009, 2012; Turkeltaub et al., 2002, 2012) implemented as in-house MATLAB tools. This algorithm aims to identify areas with across-experiment convergence of activity foci that is higher than expected from random spatial association. Before analysis, any coordinates reported in Talairach space were transformed into MNI space (Lancaster et al., 2007). Because the standard brain templates used in SPM (statistical parametric mapping) since version SPM96 and in FSL (FMRIB Software Library) are given in MNI space, reported results from analyses using SPM or FSL were treated as MNI coordinates unless the authors explicitly mentioned a transformation from MNI to Talairach space or the use of an alternative brain template.

In a first step, important content of the included studies was coded and recorded. In a second step, the reported coordinates of each experiment’s peak activations (“foci”) were projected on a brain template, acknowledging the spatial uncertainty associated with each coordinate by modeling Gaussian probability distributions around each focus. Third, the probability distributions of all activation foci were combined for each voxel, resulting in a modeled activation map. The union of these modeled activation maps then yielded voxel-wise ALE scores, which were compared to a null distribution reflecting a random spatial association between experiments. The *p*-value of a “true” ALE score was then given by the proportion of equal or higher values obtained under the null distribution. The resulting non-parametric *p* values for each meta-analysis were cut off at a threshold of *p* < .05 (family-wise error corrected at cluster level; cluster inclusion threshold at voxel level: *p* < .001).

##### 2.1.2.2. Meta-Analyses Conducted

First, a meta-analysis pooling across within-group task effects (i.e., main task effects for both age groups) was conducted on all experiments to examine the main effect of performing EF tasks on brain activity independent of age. Second, for examining age-related effects, we performed three different meta-analyses of between-group contrasts: (1) pooled, (2) old > young, and (3) young > old. We also aimed to conduct separate meta-analyses for each EF subcomponent, but only for inhibition more than 17 experiments were found to be eligible. Hence, a comparison between EF subcomponents was not possible. The results of the inhibition-specific meta-analyses can be found in Tables A5 and A6.

Pooled analyses search for consistent group differences in EF-related brain activity, independent of the direction of the between-group effect. For neuroimaging findings, pooled meta-analyses may provide the best summary because the directions of group differences in individual experiments depend on how exactly these differences were calculated, which varies widely between studies: Some authors compute task versus control contrasts at the individual-subject level, which are then compared between old and young adults at group level, whereas others compute group (old versus young) by task (task versus control or baseline) interactions at the second level. As control conditions strongly vary between experiments (from resting baseline to high-level control tasks), effects of between-group activation differences in these control conditions may influence the overall direction of group differences unpredictably (Müller et al., 2017).

### 2.2 Resting-State Functional Connectivity

To further characterize EF-related brain regions consistently affected by aging (i.e., regions with significant convergence in the pooled age-related meta-analysis), we investigated their RS-FC patterns. Therefore, whole-brain RS-FC analyses were conducted. RS-fMRI images of 413 healthy adults were obtained from the publicly available enhanced Nathan Kline Institute - Rockland Sample (eNKI-RS; Nooner et al., 2012; age range = 18 – 80; mean age = 44.85; SD = 18.51; 272 females). The re-analysis of the data was approved by the local ethics committee of the Heinrich Heine University Düsseldorf. Images were obtained with a Siemens TimTrio 3T scanner using BOLD (blood-oxygen-level-dependent) contrast [gradient-echo EPI (echo planar imaging) pulse sequence, TR = 1.4 s, TE = 30 ms, flip angle = 65°, voxel size = 2.0 × 2.0 × 2.0 mm^3^, 64 slices]. 404 volumes were acquired. Participants were instructed to keep their eyes open and maintain fixation on a central dot. Physiological and movement artifacts were removed from RS data by using FIX (FMRIB’s ICA-based Xnoiseifier, version 1.061 as implemented in FSL 5.0.9; Griffanti et al., 2014; Salimi-Khorshidi et al., 2014), which decomposes the data into independent components and identifies noise components using a large number of distinct spatial and temporal features via pattern classification. Unique variance related to the identified artifactual components is then regressed from the data. Data were further preprocessed using SPM12 (Wellcome Trust Centre Neuroimaging, London) and in-house Matlab scripts. After removing the first four dummy scans of each time series, the remaining EPI volumes were then corrected for head movement by a two-pass affine registration procedure: first, images were aligned to the initial volume and, subsequently, to the mean of all volumes. The mean EPI image was then coregistered to the gray-matter probability map provided by SPM12 using normalized mutual information and keeping all EPI volumes aligned. Next, the mean EPI image of each subject was spatially normalized to MNI-152 space using the “unified segmentation” approach (Ashburner & Friston, 2000). The resulting deformation parameters were then applied to all other EPI volumes. Finally, data were spatially smoothed with a 5-mm FWHM (full width at half maximum) Gaussian kernel to improve the signal-to-noise ratio and to compensate for residual anatomic variations.

The BOLD signal time-courses of all voxels within each seed region, expressed as the first eigenvariate, were extracted for each subject. To reduce spurious correlations, variance explained by the mean white-matter and cerebrospinal-fluid signal were removed from the time series together with 24 movement parameters (including derivatives and 2^nd^-order effects; cf. Satterthwaite et al., 2013), which was subsequently band-pass filtered with the cut-off frequencies of .01 and .08 Hz. Linear (Pearson) correlations between the time series of the seed regions and all other grey-matter voxels in the brain were computed to quantify RS-FC. The resulting voxel-wise correlation coefficients were then transformed into Fisher’s Z-scores and entered in a group-level analysis of variance. The results of this random-effects analysis were masked with the subjects’ mean Z-scores >= .1 and thresholded at a voxel-level FWE-corrected threshold of one-sided *p* < .05. Here, we chose one-sided testing, as our hypotheses were directed (i.e., we were only interested in the positive coupling between our seed regions and the rest of the brain). An additional extent threshold of 10 contiguous voxels was applied to exclude smaller, potentially spurious clusters.

### 2.3 Association of RS-FC with Age and EF Abilities

In the same sample of 413 adults, we also examined the association of the seed region’s whole-brain RS-FC with age as well as EF abilities using analysis of covariance (ANCOVA). For assessing EF abilities, we computed four compound scores: a total score and three subscores, each representing a particular EF subcomponent (i.e., working memory, inhibitory control, and cognitive flexibility). The cognitive tasks used were also obtained from the eNKI-RS. Performance raw scores were z-transformed - outliers > |3| standard deviations were removed - and added up to calculate EF subcomponent scores as follows: The working memory compound score consisted of reaction time (RT) and error rate (ER) of the 2-back and 1-back conditions of the Short Letter-N-Back Test, which is part of Penn’s Computerized Neurocognitive Battery (CNB; Gur et al., 2010). The inhibition compound score consisted of (i) the conflict effect of the Attention Network Task (ANT; Fan et al., 2002), (ii) RT and ER of the Color-Word Interference Test, which is part of the Delis-Kaplan Executive Function System (D-KEFS; Delis et al., 2004), and (iii) RT and ER of the Short Penn Continuous Performance Test (Number and Letter Versions), which is also part of Penn’s CNB. The cognitive flexibility compound score consisted of RT of the Trail Making Test, which is part of the D-KEFS, as well as of RT and ER of the Penn Conditional Exclusion Task, part of Penn’s CNB. Finally, for the total EF score, all single scores were added up and divided by the absolute number of scores. All compound scores were multiplied by -1. Hence, higher scores represent higher performance. The results of the ANCOVA were masked with the RS-FC map of the respective seed region (as described above, see section 2.2.) and thresholded at a voxel-level FWE-corrected threshold of two-sided *p* < .00625 (additional extent threshold of 10 contiguous voxels). The *p*-value was adjusted for multiple comparisons as we tested four models, and adjusted for two-sided testing as recommended by Chen et al. (2018). All results were anatomically labeled by reference to probabilistic cytoarchitectonic maps of the human brain using the SPM Anatomy Toolbox version 3 (Eickhoff et al., 2005, 2007) and visualized with the BrainNet Viewer (Xia et al., 2013).

## 3. Results

### 3.1 Meta-Analyses

#### 3.1.1. Analysis of EF-related Effects Across Age

A meta-analysis across both age groups and all experiments (reflecting all three EF subcomponents) was conducted to examine the general main effect of taxing EFs on regional brain activity. Significant convergence across experiments was found in left IFJ, left pre-SMA, left IPS/SPL, left mid-FG, left central Ins, and right frontal pole/MFG (see Table 1 and Figure 2).

**Table 1.**
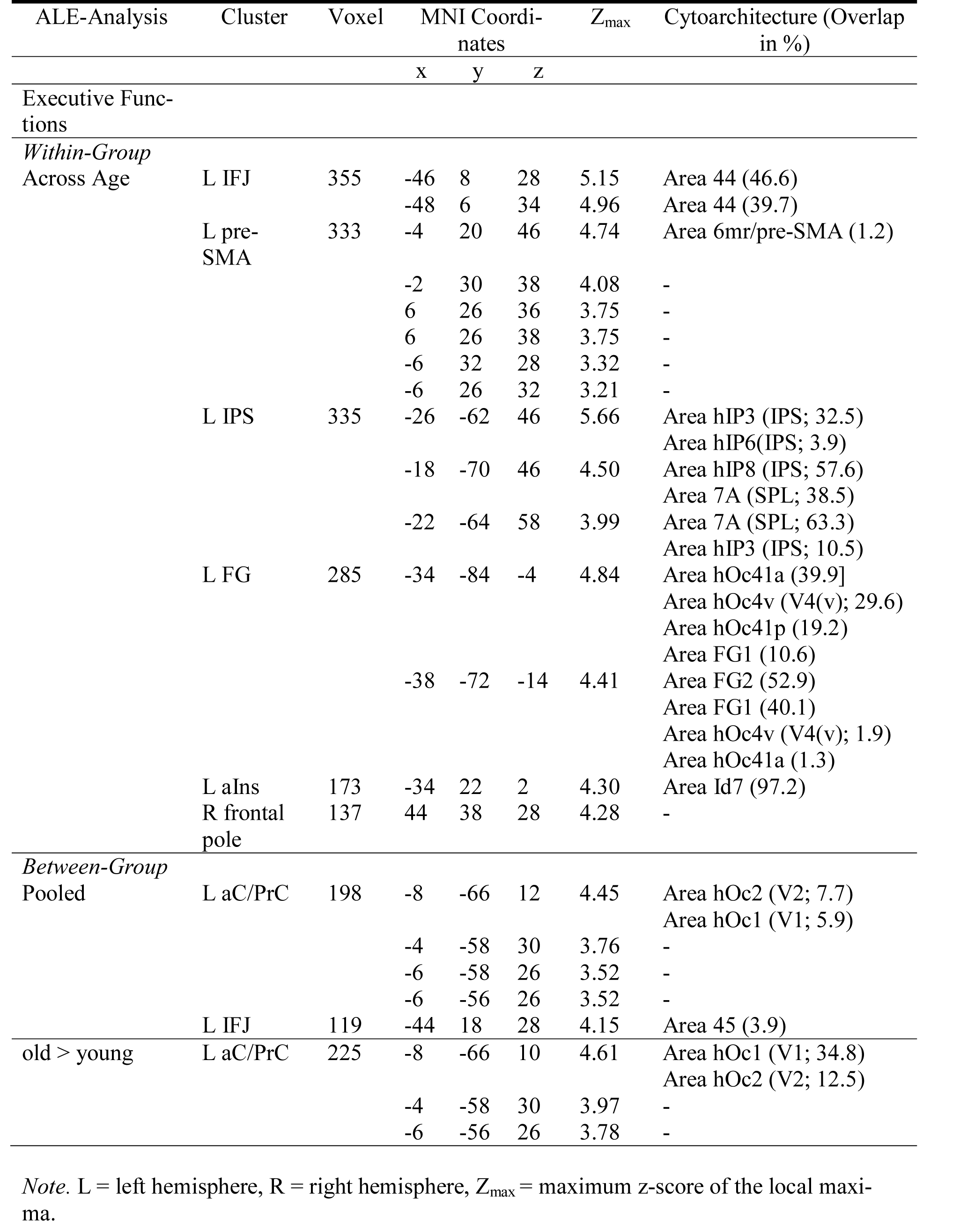
Brain Regions Showing Significant Convergence of Activity in Executive Functions

**Figure 2.**
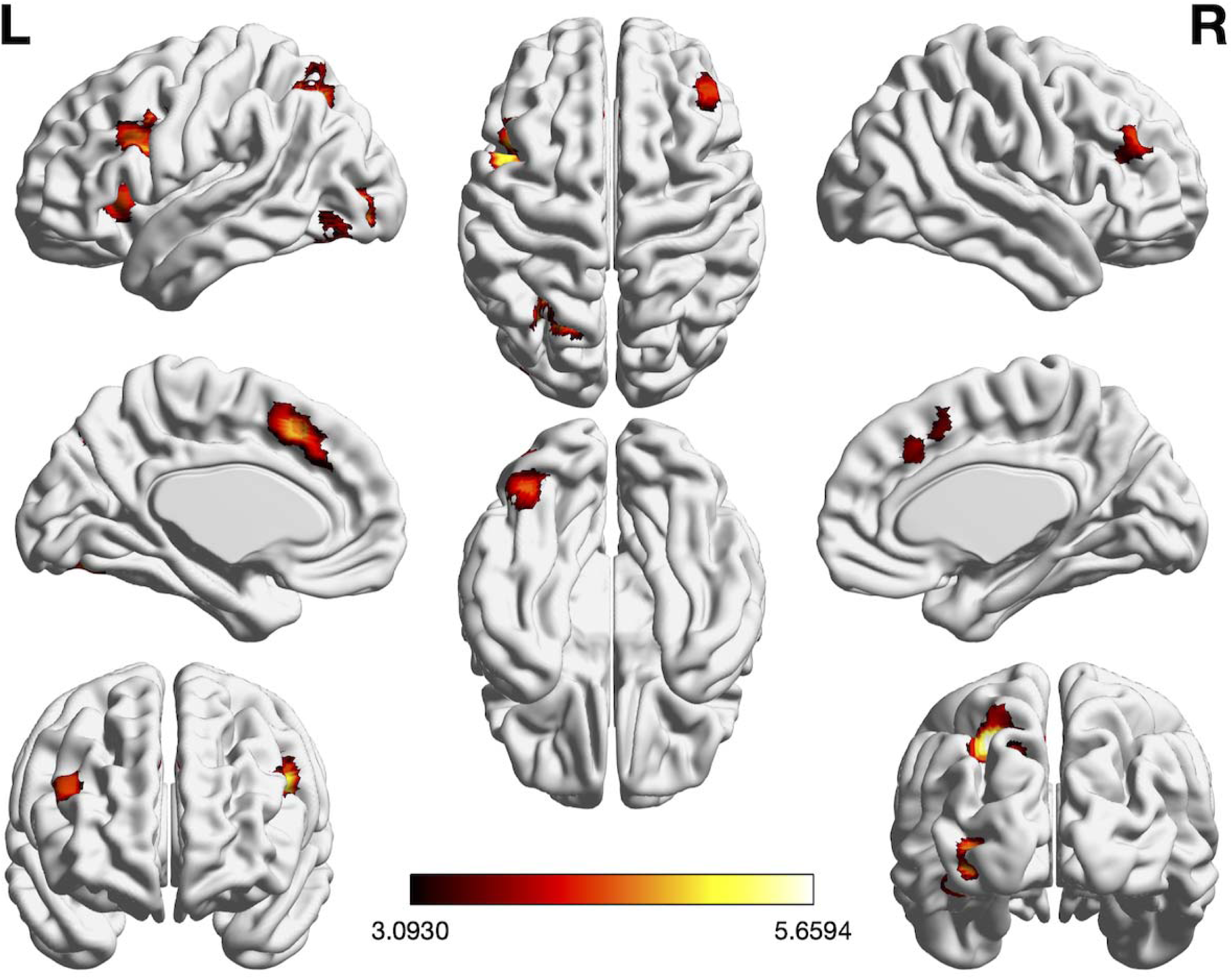
Foci of brain activity showing significant convergence of activity for EFs across age (cluster-level *p* < .05, family-wise error-corrected for multiple comparisons, cluster-forming threshold at voxel level: *p* < .001). The scale bar reflects the maximum z-score of the local maxima.

#### 3.1.2. Analyses of Age-related Differences

We performed three different meta-analyses of contrasts between age groups: (1) pooled, (2) old > young, and (3) young > old. The pooled meta-analysis, which included all experiments (n = 49) that probed age differences in EF-related brain activity irrespective of the contrast’s direction, revealed only two regions with significant convergence of such age differences: left IFJ and left anterior cuneus/precuneus (aC/PrC; see Table 1 and Figure 3A). Convergence in left IFJ was almost equally driven by experiments probing working memory (32.95%), inhibition (28.54%), and cognitive flexibility (38.41%). Furthermore, it was more strongly driven by experiments contrasting old > young (60.24%) than by experiments contrasting young > old (39.7%). Convergent activity in left aC/PrC was also driven by experiments on working memory (24.9%), inhibition (41.91%), and cognitive flexibility (32.45%). In contrast to left IFJ, however, it was almost exclusively driven by old > young contrasts (91.68%). Please see Table A4 for a full overview of the study contributions.

**Figure 3.**
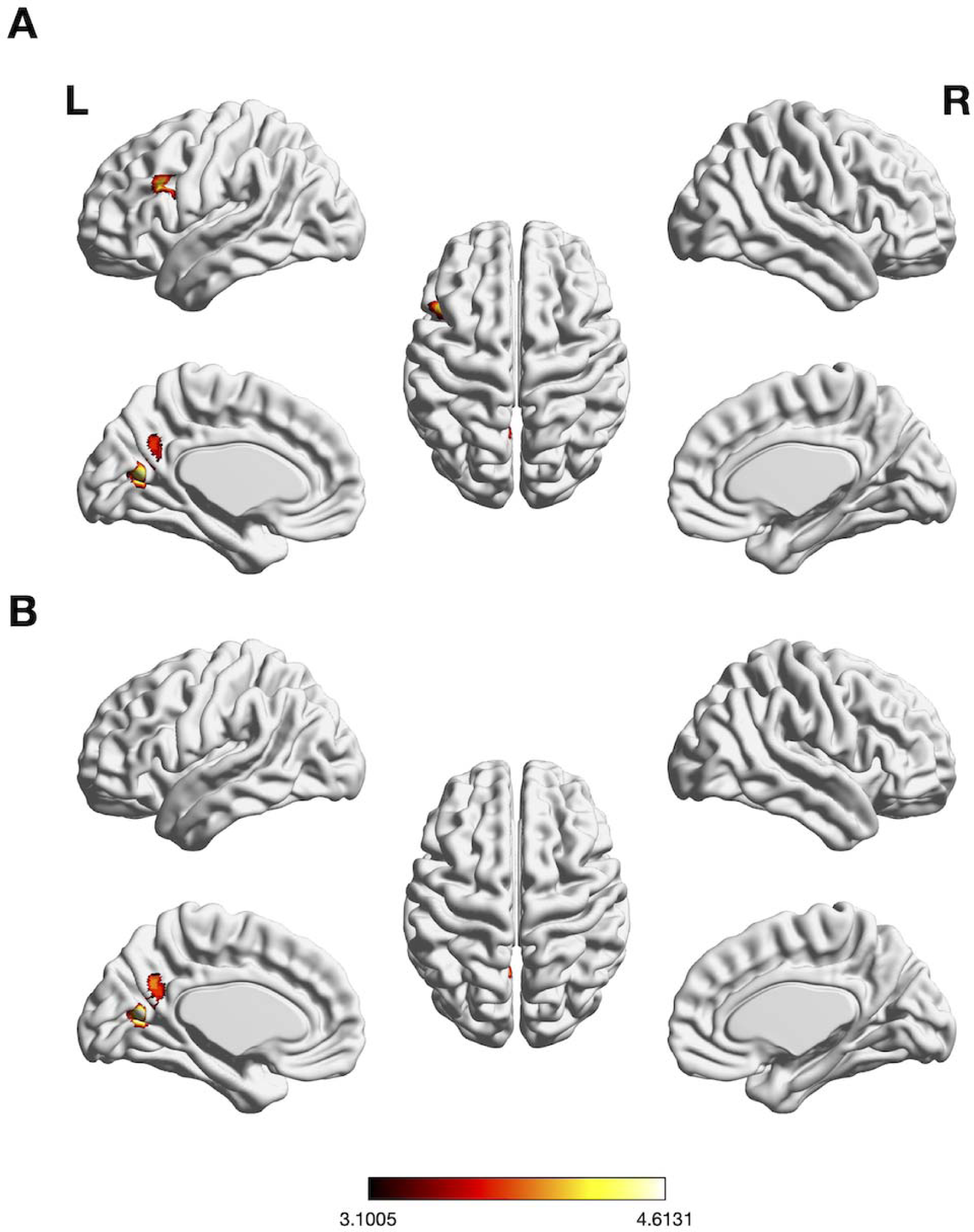
Foci of brain activity showing significant convergence of activity for (A) EFs pooled, (B) EFs old > young (cluster-level *p* < .05, family-wise error-corrected for multiple comparisons, cluster-forming threshold at voxel level: *p* < .001). The scale bar reflects the maximum z-score of the local maxima.

The meta-analysis testing for consistently lower brain activity across EF experiments in older (vs. younger) adults (n = 31) did not yield any significant convergence. Conversely, the meta-analysis testing for consistently higher activity across EFs experiments in older (vs. younger) adults (n = 42) revealed significant convergence in left aC/PrC (see Table 1 and Figure 3B).

We also aimed to conduct separate meta-analyses for each EF subcomponent, but only for inhibition more than 17 experiments were found to be eligible. The results of the inhibition-specific meta-analyses can be found in Tables A5 and A6.

### 3.2 Connectional Characterization

The two age-sensitive regions resulting from the pooled meta-analysis (i.e., left IFJ and left aC/PrC) were connectionally characterized by conducting whole-brain RS-FC analyses. The RS-FC map obtained for left IFJ comprised 14 clusters of significant coupling: the seed region extending into DLPFC, MFG, FEF, dPMC, SMA/pre-SMA, frontal pole, and aIns; left caudate nucleus; left IPS extending into FG and SPL; two clusters in left cerebellum VII; right IFJ extending into DLPFC, FEF, dPMC, and frontal pole; right cerebellum VI and VII; right IPS/angular gyrus; right FG extending into Wernicke’s region; right SMA/pre-SMA; right aIns, right primary somatosensory cortex (S1); and bilateral anterior cingulate cortex (ACC; see Table 2 and Figure 4A).

**Figure 4.**
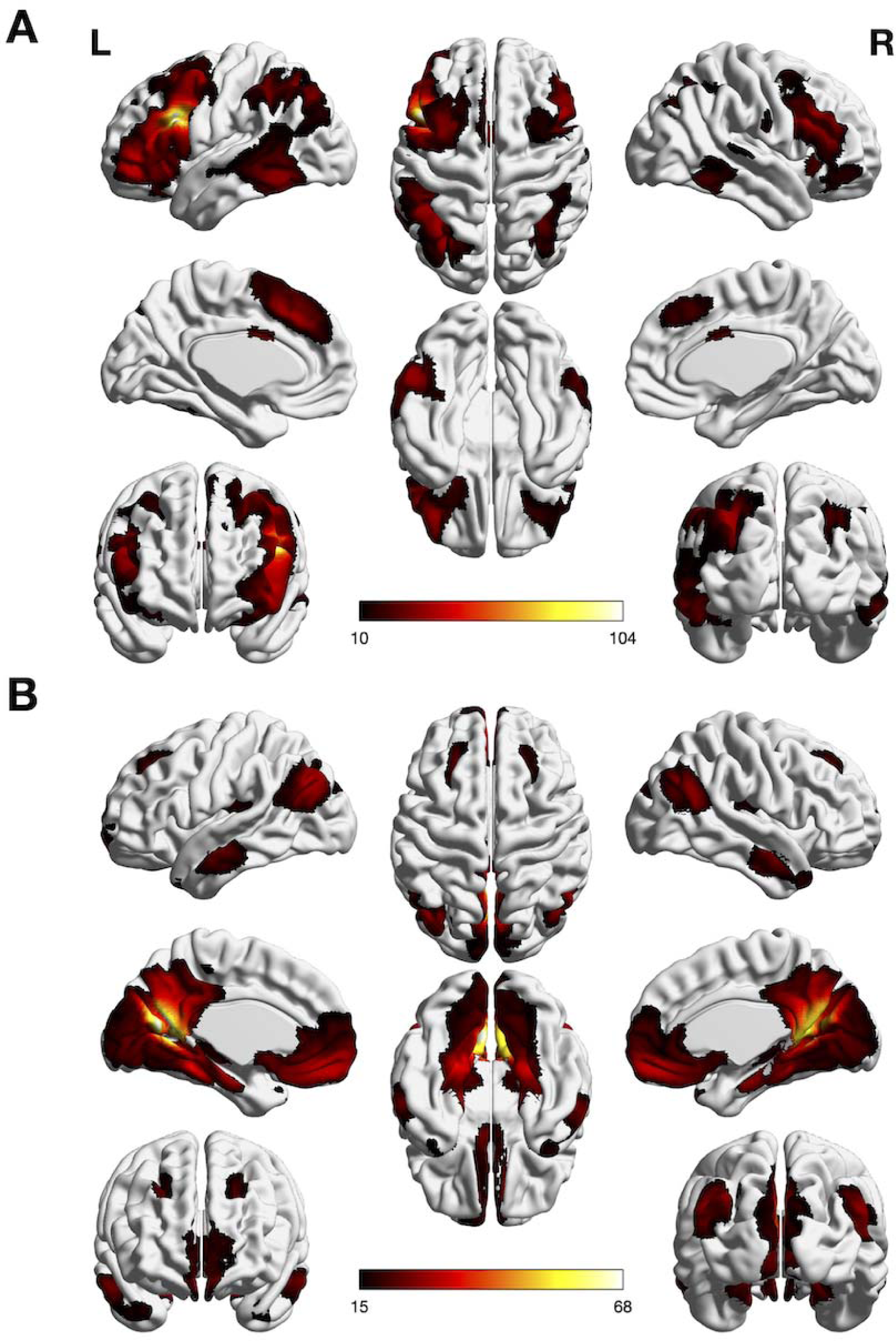
Whole-brain RS-FC analyses of (A) left IFJ and (B) left aC/PrC (voxel-level family-wise error corrected threshold of one-sided *p* < .05, extent threshold = 20, masked with the subjects’ mean Z-scores >=.I). The scale bar reflects t-scores.

**Table 2.**
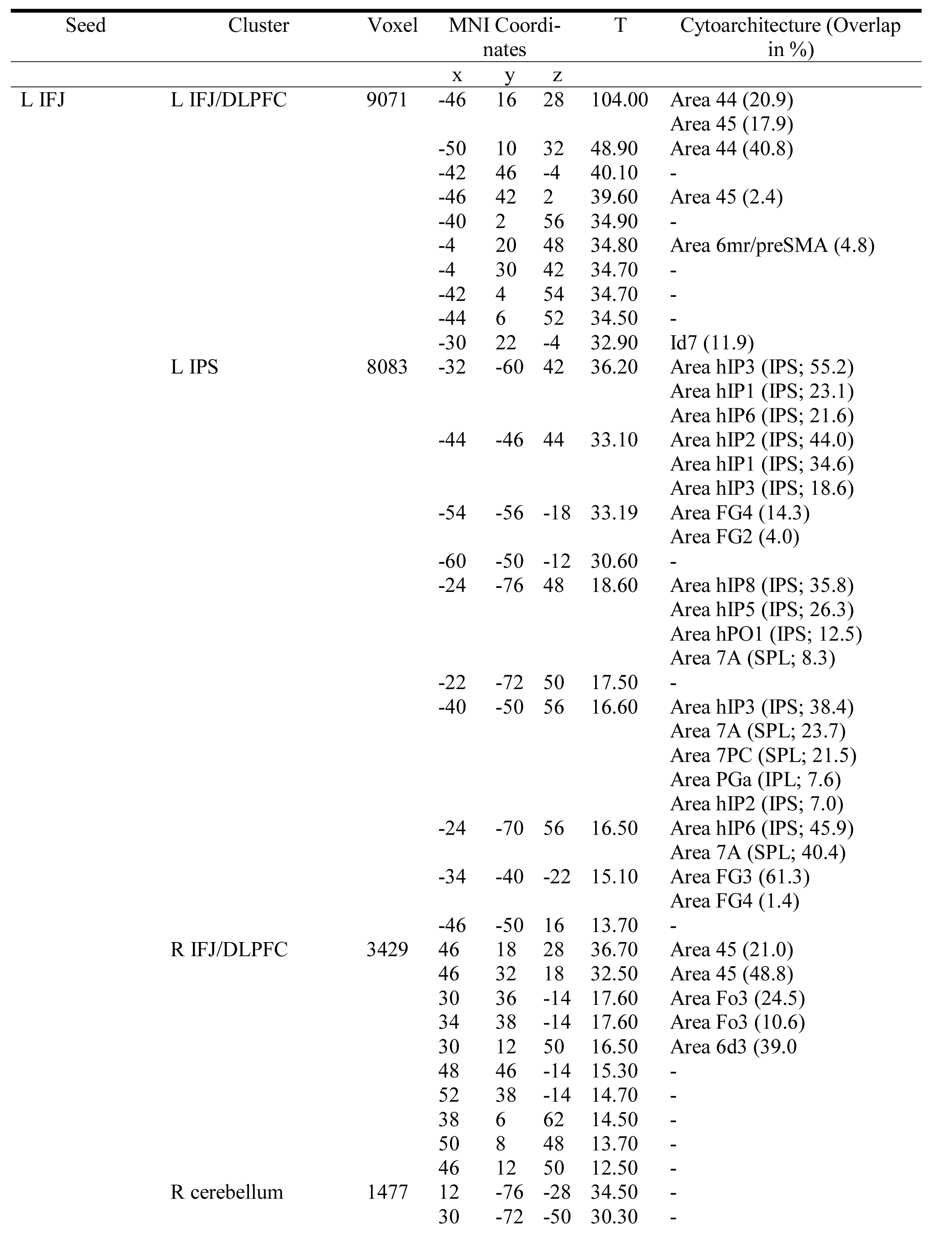
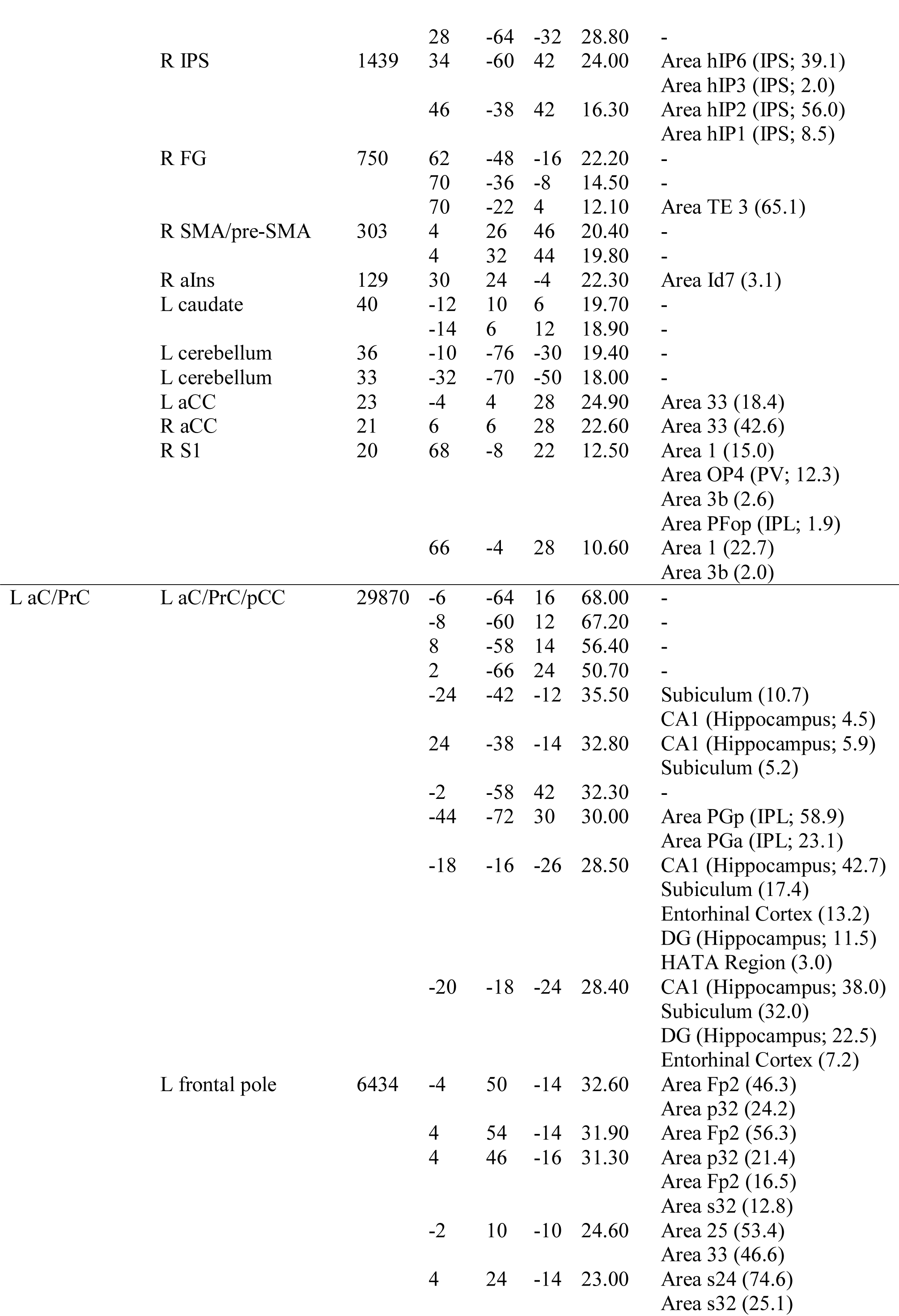
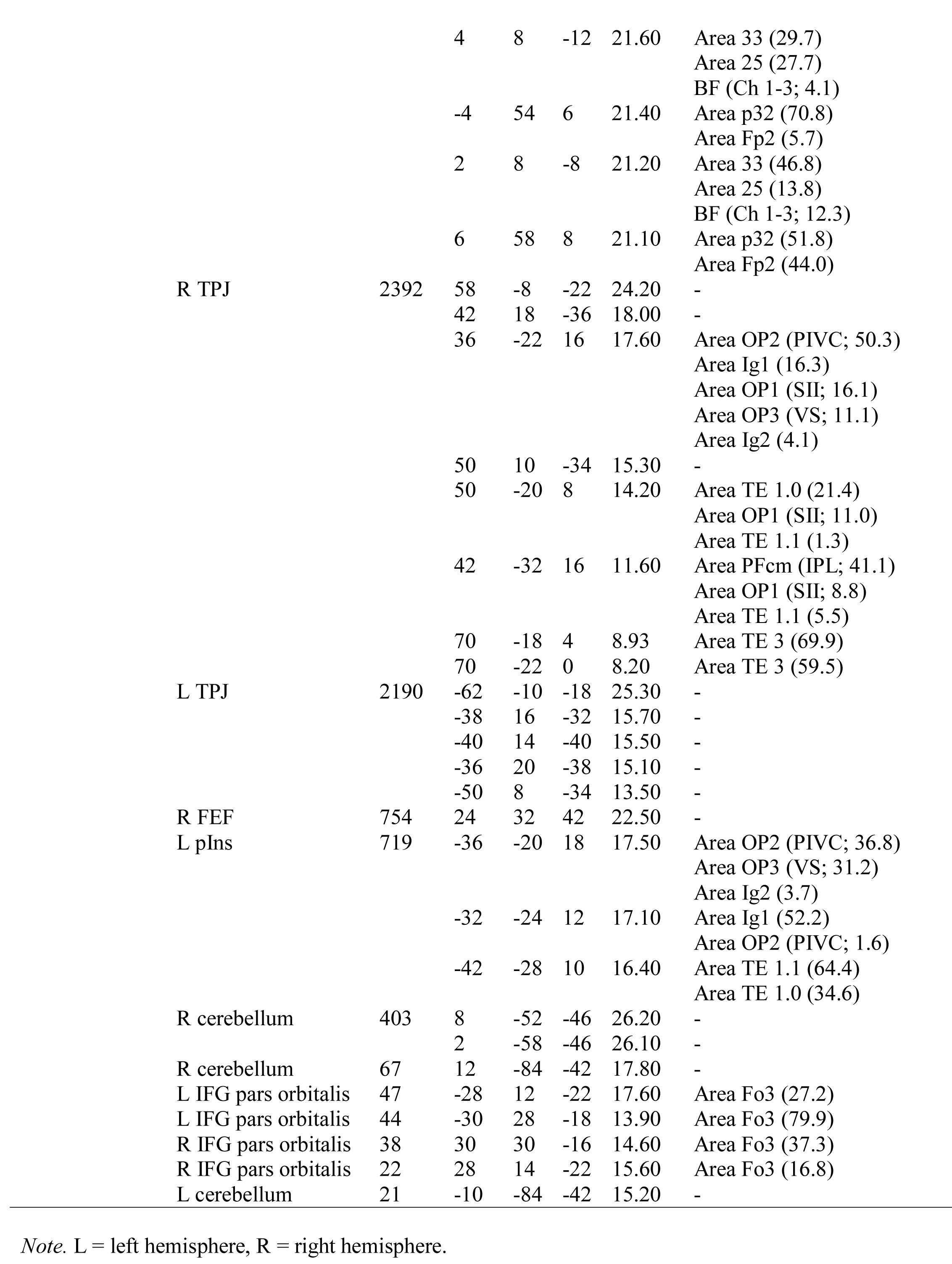
RS-FC Analyses

The RS-FC analysis of left aC/PrC yielded 13 clusters: the seed region extending into bilateral PCC, FG, subiculum, calcarine gyrus, and left IPL; left frontal pole extending into subgenual area, FEF, and bilateral frontopolar cortex; left posterior Insula (pIns) extending into parietal operculum; left cerebellum VII; two clusters in right cerebellum IX and VII; bilateral temporoparietal junction (TPJ); and four clusters in bilateral IFG pars orbitalis (see Table 2 and Figure 4B).

### 3.3 Association of RS-FC with Age and EF Abilities

#### 3.3.1. Age

An ANCOVA was performed to examine the association between the seed regions’ RS-FC patterns and age. We observed significant negative associations with age for RS-FC between left IFJ and 10 clusters: left aIns, left FEF, left TPJ, bilateral IFJ/DLPFC, bilateral FG, and bilateral aCC (see Table 3 and Figure 5A).

**Figure 5.**
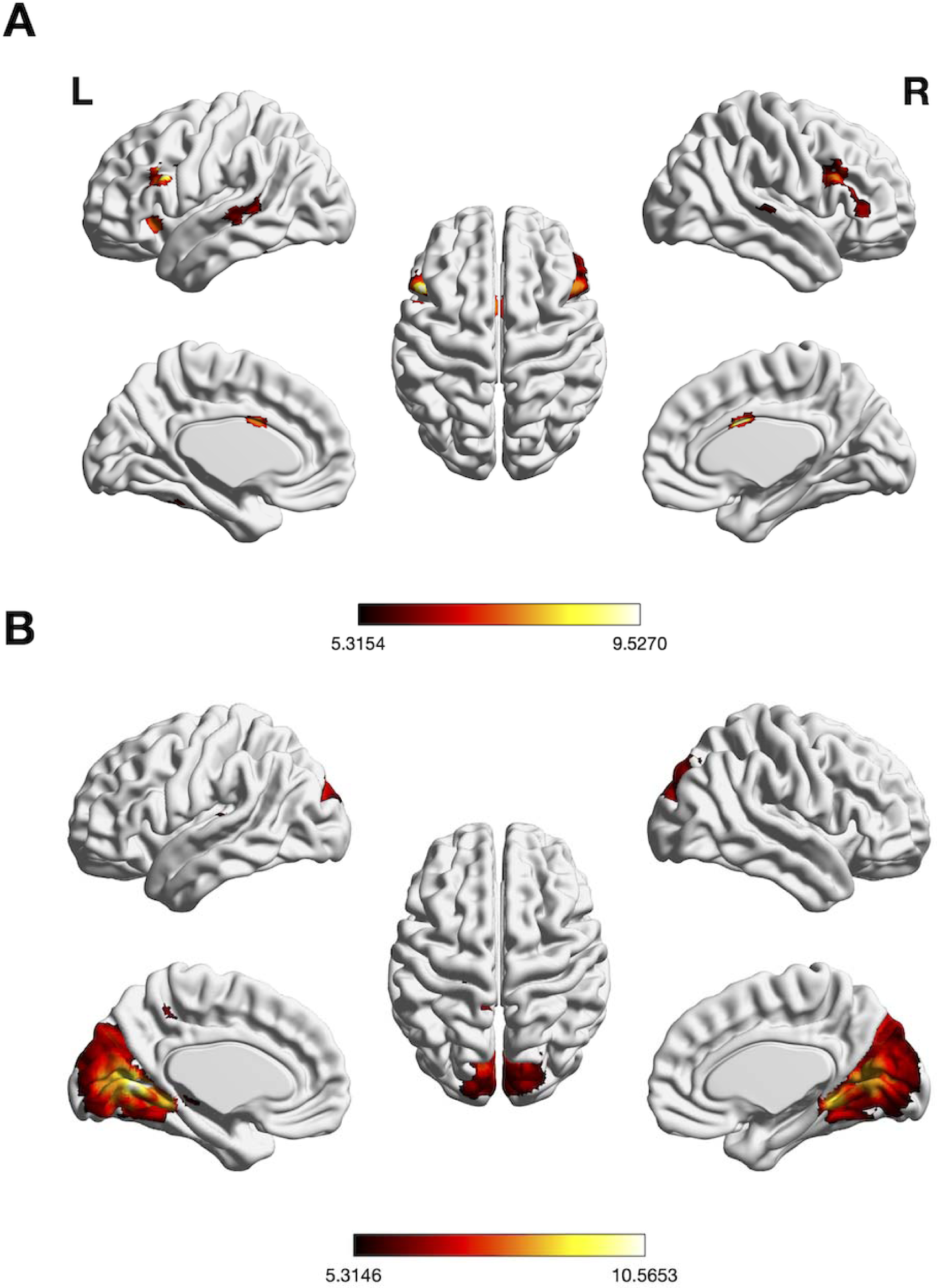
Significant negative association between whole-brain RS-FC of (A) left IFJ and age and (B) aC/PrC and age, (voxel-level family-wise error-corrected threshold of two-sided *p* <.00625, extent threshold = 10, masked with RS-FC map of left IFJ and aC/PrC, respectively). The scale bar reflects t-scores.

**Table 3.**
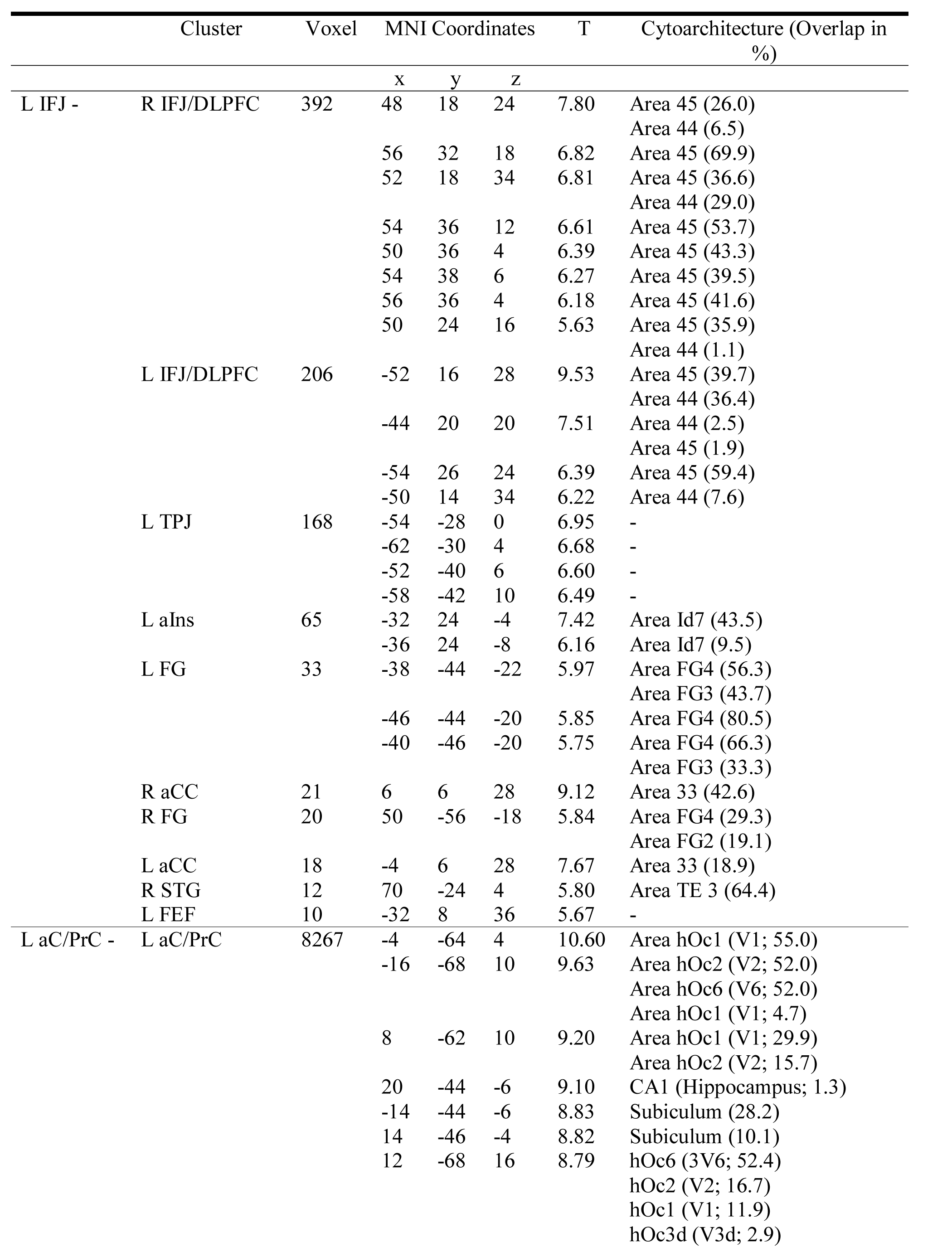
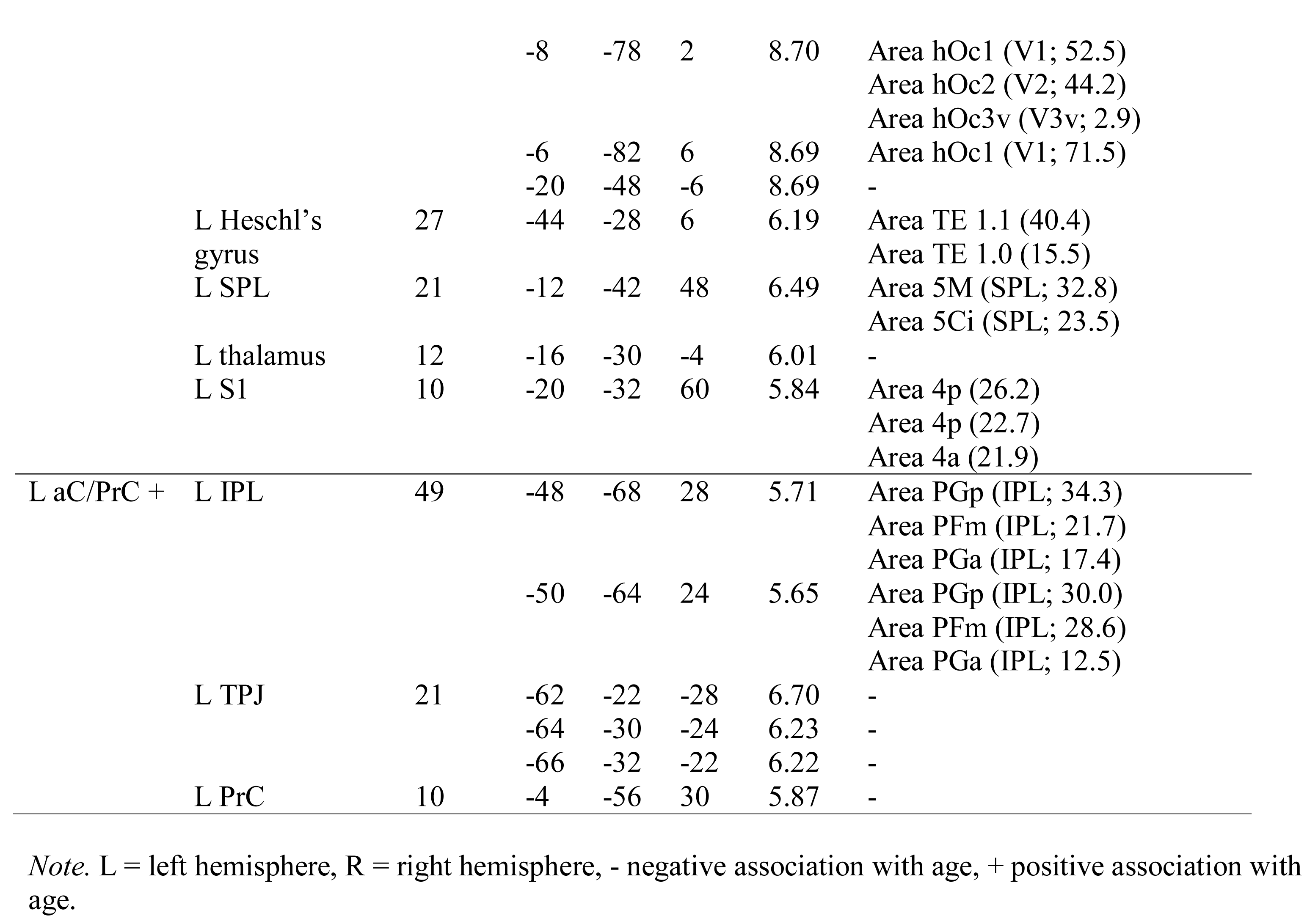
Association of RS-FC and Age

Age was also significantly negatively associated with RS-FC between left aC/PrC and 5 regions: the seed region extending into bilateral visual cortex, left Heschl’s gyrus extending into planum temporale, left SPL, left S1, and left thalamus (see Table 3 and Figure 5B). Finally, age was significantly positively associated with RS-FC between left aC/PrC and 3 regions: left IPL, left PrC, and left TPJ (see Table 3 and Figure 6).

**Figure 6.**
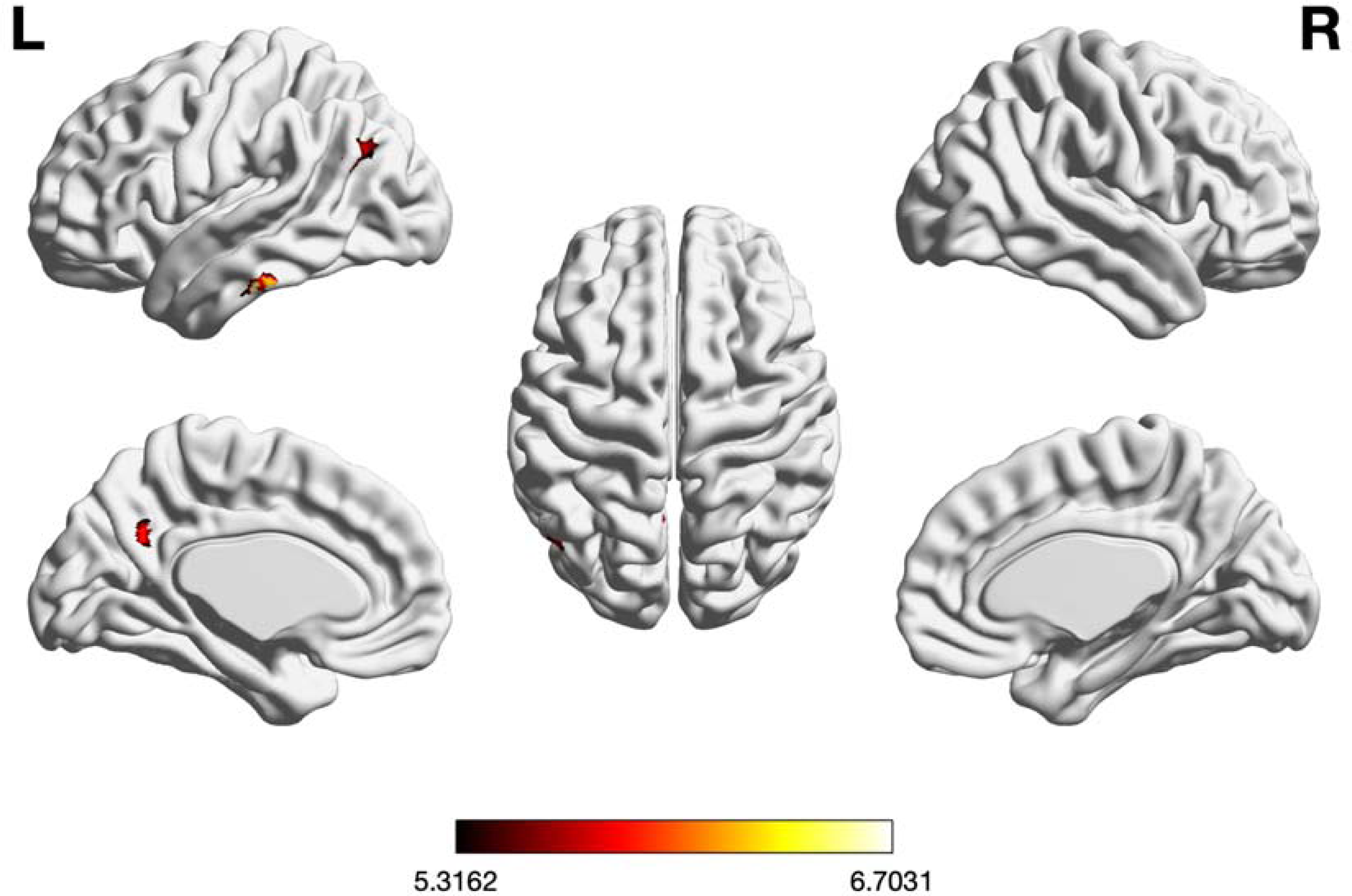
Significant positive association between whole-brain RS-FC of left aC/PrC and age, (voxel-level family-wise error-corrected threshold of two-sided *p* < .00625, extent threshold = 20, masked with RS-FC map of left aC/PrC). The scale bar reflects t-scores.

#### 3.3.2. EF Abilities

Finally, we performed an ANCOVA to assess the association between the seed regions’ RS-FC patterns and EF abilities. RS-FC between left aC/PrC and the seed region extending into bilateral visual cortices was significantly positively associated with the total EF score (see Table 4 and Figure 7).

**Figure 7.**
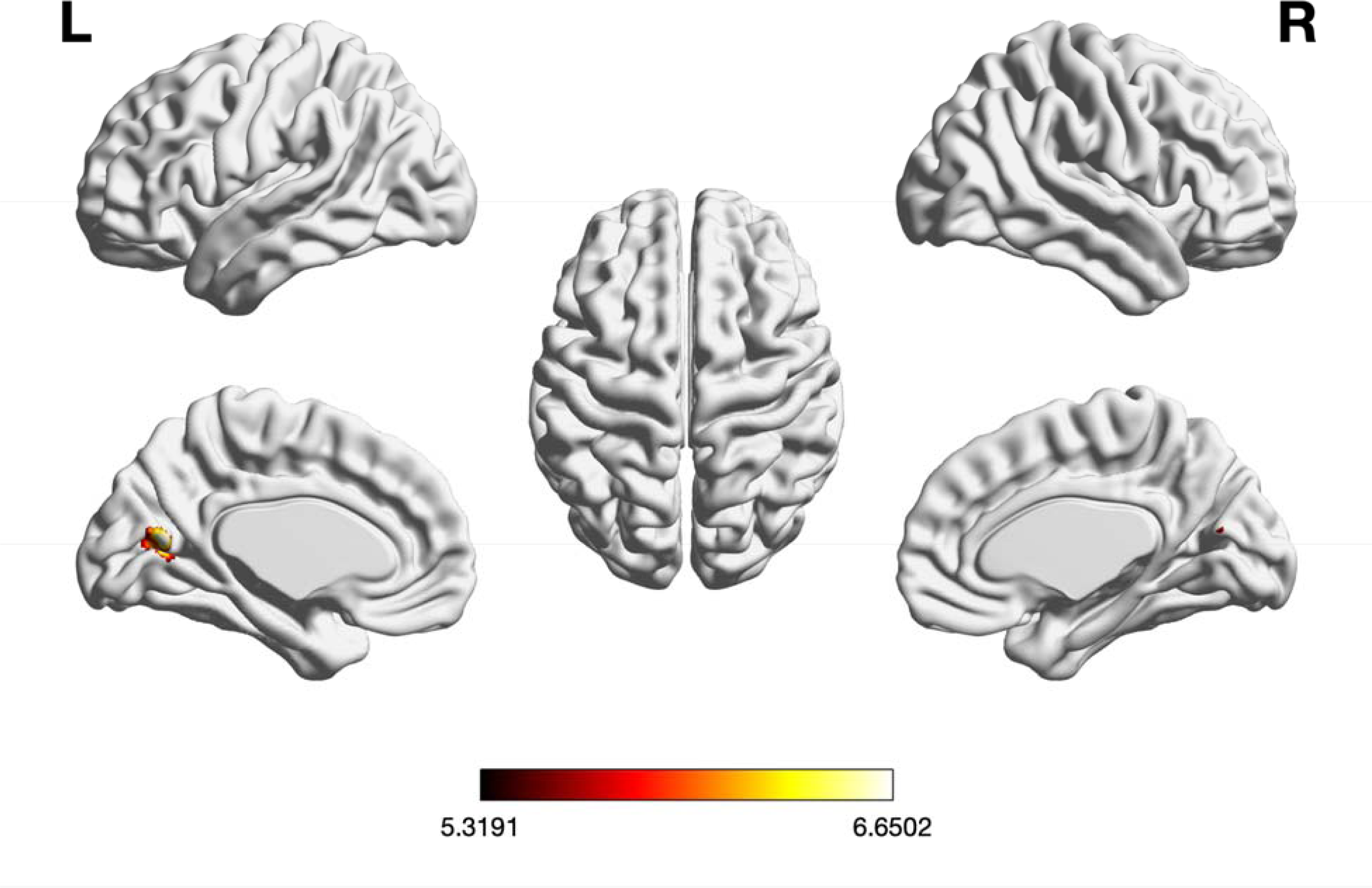
Significant positive association between whole-brain RS-FC of left aC/PrC and executive functions, (voxel-level family-wise error-corrected threshold at two-sided *p* < .00625, extent threshold = 10, masked with RS-FC map of left aC/PrC). The scale bar reflects t-scores.

**Table 4.**
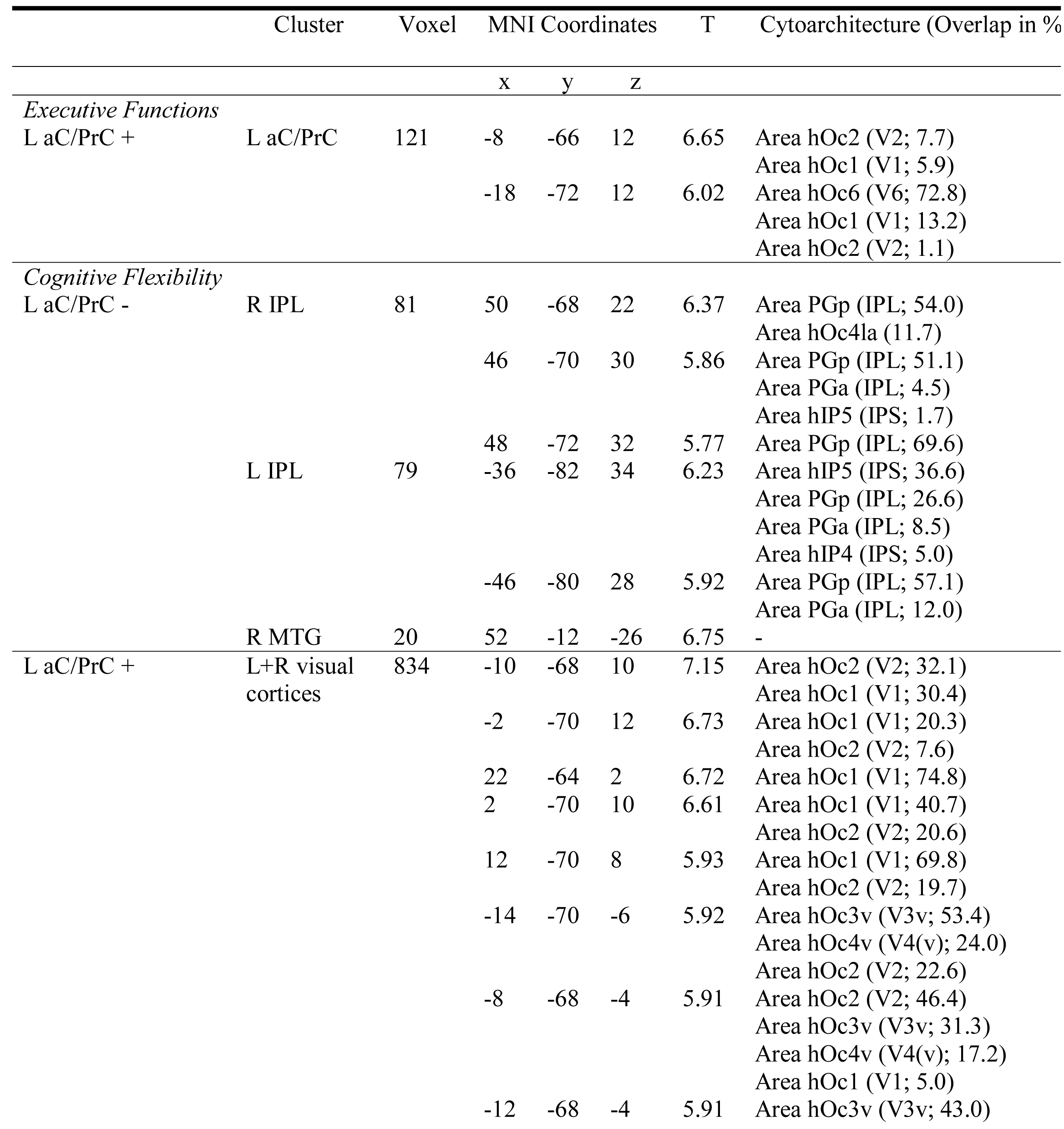
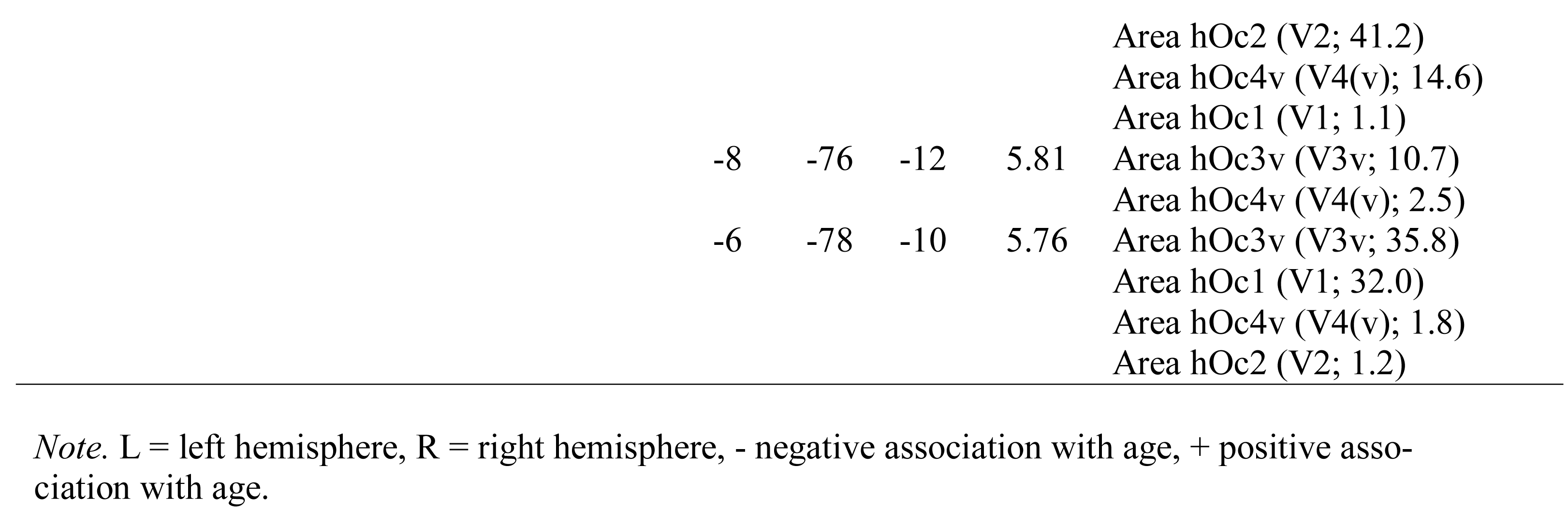
Association of RS-FC and Combined Total Executive Functions and Cognitive Flexibility Compound Scores

We found a significant negative association of the cognitive flexibility score with RS-FC between left aC/PrC and 3 regions: bilateral IPL and right middle temporal gyrus (MTG; see Table 4 and Figure 8A), whereas RS-FC between left aC/PrC and the seed region extending into bilateral visual cortices was significantly positively associated with the cognitive flexibility score (see Table 4 and Figure 8B).

**Figure 8.**
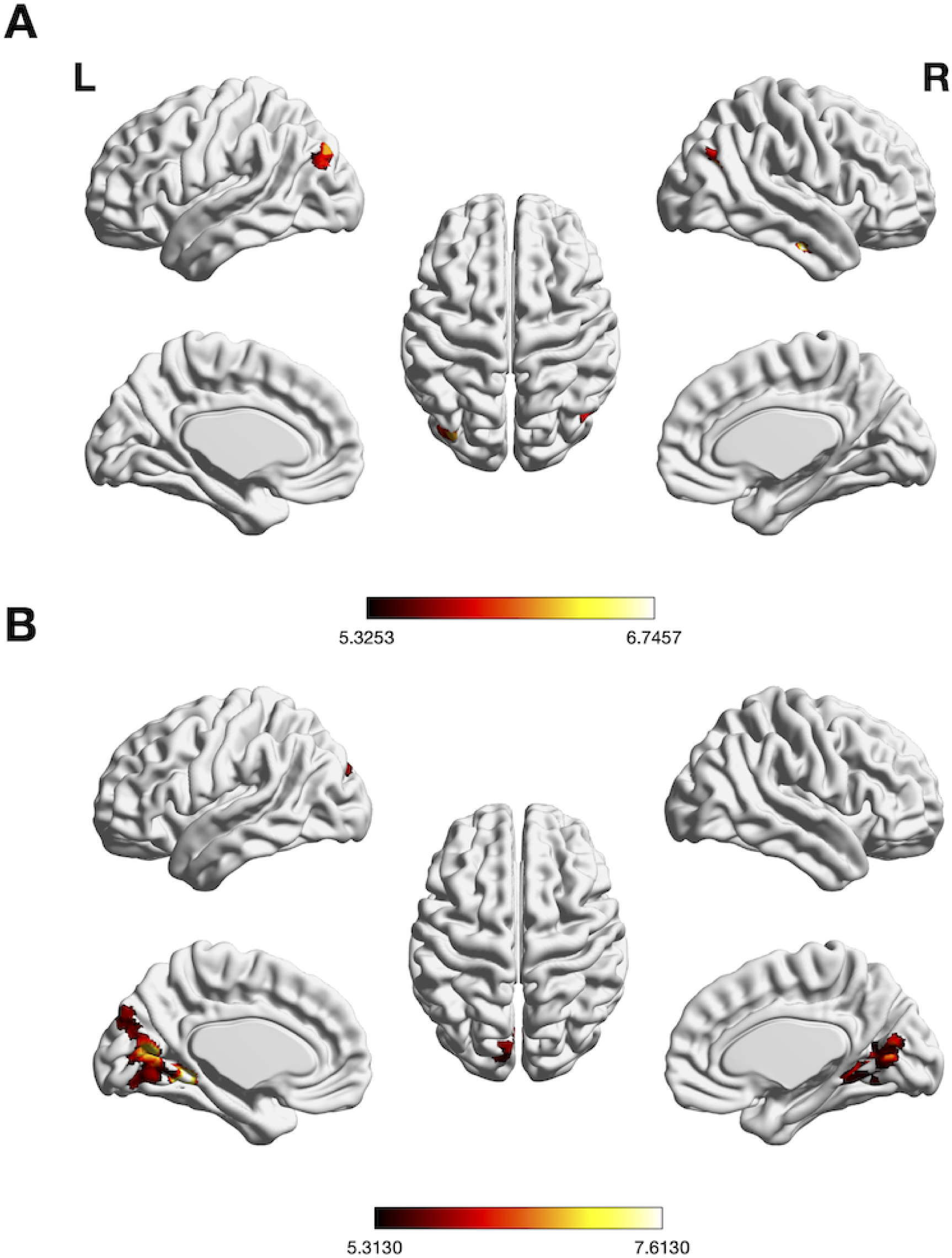
(A) Significant negative and (B) positive association between whole-brain RS-FC of left aC/PrC and cognitive flexibility, (voxel-level family-wise error-corrected threshold at two sided *p* < .00625, extent threshold = 10, masked with RS-FC map of left aC/PrC). The scale bar reflects t-scores.

Neither for working memory nor for inhibitory control was there any significant association between performance (compound scores) and RS-FC of either seed region with the rest of the brain.

## 4. Discussion

Coordinate-based ALE meta-analyses were used to synthesize the neural correlates of age-related changes in EFs. In particular, we first ran a meta-analysis across all age groups and all three EF subcomponents followed by a pooled and two directed meta-analyses examining age differences in EF-related brain activity. The initial global analysis corroborated a set of regions well-known for being involved in EFs. Consistent activation differences between young and old adults, however, were restricted to left IFJ and left aC/PrC. Subsequently, we assessed the connectional profiles of these two age-sensitive regions and how their RS-FC profiles are modulated by age and EF abilities. Left IFJ was found to be linked to regions involved in executive functioning, whereas left aC/PrC was connectionally linked to regions involved in attentional processes and the DMN. Furthermore, RS-FC between left IFJ and EF-related regions decreased with increasing age. Similarly, RS-FC between left aC/PrC and regions involved in perceptual processes decreased with increasing age, while RS-FC between left aC/PrC and DMN-related regions increased with age. Finally, only very few associations of seed-based RS-FC with EF abilities were observed: RS-FC between left aC/PrC and bilateral visual cortex was positively associated with the total EF score and cognitive flexibility, whereas RS-FC between left aC/PrC and DMN-related regions was inversely associated with cognitive flexibility.

### 4.1 Comparison to Previous Meta-Analyses

The current results of between-group contrasts deviate quite noticeably from previous meta-analyses of age differences in EF-related brain activity (Di et al., 2014; Spreng et al., 2010; Turner & Spreng, 2012). The only two regions consistently found across these earlier meta-analyses are left IFJ and pre-SMA. Thus, there is substantial disagreement between all meta-analyses devoted to this topic.

These discrepancies might be explained by several methodological differences: First, all previous meta-analyses included several reports of ROI analyses, which biases ALE whole-brain tests for significance towards convergence in the given ROIs (Müller et al., 2017). The null distribution in ALE reflects a random spatial association between findings across the entire brain assuming that each voxel has the same a priori chance of being activated (Eickhoff et al., 2012). The inclusion of ROI analyses would obviously violate this assumption, leading to inflated significance estimations for regions supported by ROI analyses (Müller et al., 2017). Second, all previous meta-analyses attempted to correct for multiple comparisons by controlling the voxel-level FDR, which is considered invalid for topographic inference on smoothed data (Chumbley & Friston, 2009), features low sensitivity, and leads to inflated positive findings (Eickhoff et al., 2016). FWE correction for ALE meta-analyses on the other hand provides good sensitivity and low susceptibility to false positives. Third, previous meta-analyses were partly based on rather small samples, rendering them prone to yielding clusters of “convergence” driven by very few or even single experiments (Eickhoff et al., 2016). Fourth, earlier analyses included some tasks that, according to our definition, would not constitute clear-cut operationalizations of EFs (e.g., sentence comprehension or word generation tasks). Taken together, the inclusion of ROI studies, heterogeneity in the tasks included, limited sample sizes, and FDR-corrected thresholding may have rendered previous meta-analyses very liberal, leading to more widespread but potentially spurious convergence across published results.

### 4.2 Left IFJ

The pooled meta-analysis of age differences in EF-related brain activity yielded convergence in left IFJ. Our data indicate that left IFJ is recruited to a different degree by younger versus older adults. The sign of this difference, however, appears to depend on the type of task: For tasks taxing working memory, many studies report an age-related decrease in IFJ activation (e.g., Bäckman et al., 2011; Podell et al., 2012; Prakash et al., 2012). Podell et al. (2012) argued that deficits in working memory updating in older adults are accompanied by a reduced utilization of efficient neurocognitive strategies, relative to younger adults. This is in line with the dedifferentiation hypothesis of cognitive aging, stating that brain regions showing specialized responses to specific cognitive tasks become less specialized with increasing age (Baltes & Lindenberger, 1997; Goh, 2011; Li & Sikström, 2002; Park et al., 2001; 2004). In the context of inhibitory control and attention shifting, however, studies report an age-related increase in left IFJ activity (e.g., Korsch et al., 2014; Townsend et al., 2006; Zysset et al., 2007). According to Townsend et al. (2006), the more extensive activation patterns observed in older adults may be due to (i) the failure of within-channel inhibition of irrelevant visual information, or (ii) compensatory neural recruitment caused by the attempt to increase relevant and decrease irrelevant information processing. This is in line with Korsch et al.’s (2014) conclusion that increased age-related IFJ activation is caused by the use of different strategies when irrelevant information interferes with correct response selection. Looking at the individual study contributions to our cluster, our results support these findings. For experiments on cognitive flexibility or inhibition that contributed to the cluster, convergence in left IFJ was mainly driven by the contrast old > young (rather than young > old). In contradistinction, for experiments on working memory, convergence was mainly driven by the contrast young > old (rather than old > young; see Table A4). These findings, although purely descriptive, point to a shared cognitive mechanism in the context of inhibition and cognitive flexibility, possibly leading to the observed similar aging effects on IFJ activity.

In the literature, there also is a well-established link between left IFJ and task switching, set shifting, or updating task representations (Brass & Cramon, 2004; Derrfuss et al., 2005; Worringer et al., 2019), that is, processes that allow adjusting behavior to new external demands in a top-down fashion (i.e., cognitive flexibility). This notion is also supported by repetitive transcranial magnetic stimulation studies (Higo et al., 2011; Zanto et al., 2011), pointing to IFJ’s causal participation in updating task representations and regulating neural excitability in visual areas according to the task goal. Supporting the broad involvement of left IFJ across EF domains, Derrfuss et al. (2004) mapped the activity from experiments investigating working memory, task switching, and inhibitory control and found a significant overlap in IFJ for all task types. The almost equal contribution of working memory, inhibition, and cognitive flexibility experiments to the IFJ cluster in the pooled EF meta-analysis also points to its importance for all EF subcomponents. Further indirect evidence is provided by IFJ’s location at the junction of the inferior frontal and inferior precentral sulci, and thus at the intersection of three functional neuroanatomical domains: premotor, language, and working memory. Although our study cannot clarify the precise functional role of left IFJ, this region may integrate information from these three domains (Brass et al., 2005). In particular, it is thought to (re)activate and implement relevant stimulus–response mappings, connecting stimulus information with motor output according to behavioral goals (Hartstra et al., 2012; Worringer et al., 2019).

Our RS-FC results further stress left IFJ’s important role in EFs, as its RS-FC map is highly overlapping with Camilleri et al.’s (2018) eMDN, the proposed neural correlate of EFs and with the frontoparietal control network (FPCN; Cole & Schneider, 2007), that is, bilateral ACC/pre-SMA, DLPFC, IFJ, aIns, dPMC, PPC. The negative association between RS-FC of left IFJ and age (see Figure 5A) indicates that age-related connectivity changes are not regionally specific (e.g. prefrontal) but rather wide-spread, including the dorsal attention network (DAN), the FPCN as well as the eMDN. An age-related RS-FC decline in these networks has been reported previously (Campbell et al., 2012; He et al., 2014). The frequently reported age-related decline in EF performance might thus be associated with decreased FC between regions and networks important for executive functioning. Through its functional role, that is, stimulus–response mapping and its importance for all EF subcomponents, left IFJ seems to be operating as a key node for executive functioning and thus showing domain-general recruitment as well as intrinsic correlations to multiple task positive networks.

Summing up, our meta-analytic and connectional findings suggest a pivotal role of left IFJ in EFs. While its involvement in EFs may mostly be domain-general, its recruitment appears to change with age depending on the type of task. As older adults seem to rely more on left IFJ in the context of cognitive flexibility and inhibition, younger adults recruit it more strongly in the context of working memory. Decreased RS-FC with age of left IFJ and regions associated with different task positive networks points to (i) generalized age-related changes across the brain rather than degradation in a particular region, as well as (ii) a possible underlying neural correlate for EF performance decline with age.

### 4.3 Left anterior Cuneus/Precuneus

Convergence in left aC/PrC was found in the meta-analyses EF pooled and EF old > young. To account for the difficulties in accurately comparing anatomical locations across individuals and studies due to individual differences as well as differences in spatial processing and brain templates (Brett et al., 2002) we chose to label the region of convergence aC/PrC instead of deciding on just one region and thus neglecting important functional implications. Taking the contribution of our region into account, convergence in the pooled meta-analysis was mainly driven by the contrast old > young. Consequently, consistent increased activation in aC/PrC was specific to older compared to younger adults. Furthermore, it has been associated with initiating shifts of attentional focus (Bzdok et al., 2015; Langner & Eickhoff, 2013; Worringer et al., 2019). This is in accordance with our finding of activity convergence in left aC/PrC being driven by the subcomponents inhibition and cognitive flexibility, where shifting the attentional focus and thus inhibiting irrelevant input plays a key role (see Table A4). Previous studies (DiGirolamo et al., 2001; Kuptsova et al., 2016; Townsend et al., 2006) testing age-related differences in attention shifting suggest that younger and older adults relied on the same regions during shift conditions, that is, frontoparietal regions including PrC. Older adults, however, also recruited these regions during the control condition, (i.e. attentional focusing). The authors suggested that older adults relied more on executive networks, even in the non-shift task condition, to compensate for reduced efficiency of sensory and cognitive processing. Another explanation might be that older adults had difficulties inhibiting the alternate task even during the non-shift condition. By inspecting the study contributions to the left aC/PrC cluster in the pooled EF meta-analysis, one can see that 92% of the studies leading to a convergence in left aC/PrC result from the contrast old > young. 83% of these studies did not report any inclusive masking with a task-positive effect, and 68% tested against an active control condition, rather than rest. While we did not directly investigate deactivations – due to the lack of studies available that matched our inclusion criteria – one could argue, based on these numbers, that convergence in left aC/PrC might be mainly driven by consistently greater aC/PrC deactivation in older adults during the control (vs. task) condition and/or consistently greater deactivation in younger adults during the experimental (vs. control) task, rather than a higher task-induced aC/PrC activation in older adults. A greater age-related deactivation during control (vs. task) and deactivation difficulties (compared to younger adults) in task (vs. control) could lead to inefficiencies in attentional switching in older adults. Together with PCC, PrC is assumed to be one of the central and specialized hubs of the DMN, being intrinsically connected to the DMN as well as to attentional networks, in line with our RS-FC findings (see Figure 4B). Its role might be controlling the dynamic interaction between these networks for an efficient distribution of attention (Leech et al., 2011). Furthermore, PrC appears to be in a special position within the DMN as it is coupled with the DMN at rest, and with task positive networks during task performance (Leech et al., 2011; Utevsky et al., 2014). Its wide-spread FC pattern, involving higher association regions, corroborates an important role in integrating internally and externally driven stimulus processing (Cavanna & Trimble, 2006).

While PrC’s RS-FC with sensorimotor regions decreased in older adults, its RS-FC with regions associated with the DMN and DAN increased with age. Previous studies found that older adults failed to deactivate the DMN during a range of cognitive tasks (e.g., Grady et al., 2006; Lustig et al., 2003; Park et al., 2010; Persson et al., 2007). Spreng and Schacter (2011) assumed that this is due to a reduction of large-scale network flexibility in the context of changing task demands. These differences might also be due to differences during fixation, as older adults have a reduced susceptibility to mind wandering (Giambra, 1989; Jackson & Balota, 2012). Furthermore, it might be more difficult for older adults to fixate the cross, possibly explaining an age-related RS-FC increase of left PrC with the DAN. Additionally, it has been proposed that functional networks become less specific with age (Geerligs, Maurits, et al., 2014; Geerligs, Renken, et al., 2014). Thus, there might be a dedifferentiation in activation patterns – in accordance with the aforementioned dedifferentiation hypothesis of neural aging – and a compensatory recruitment of further brain regions. The latter has also been proposed by the cognitive aging theories CRUNCH (Reuter-Lorenz & Cappell, 2008) and STAC (Park & Reuter-Lorenz, 2008), which state that in older adults, to maintain cognitive and behavioral performance, connections that have become fragile or deficient are weakened, existing connections are strengthened, and new connections are developed.

RS-FC between left aC/PrC and bilateral visual cortices showed a positive association with the total EF and cognitive flexibility score, whereas RS-FC between left aC/PrC and both bilateral IPL and right MTG revealed negative associations with the latter score. While larger RS-FC of PrC and visual areas seems to support cognitive flexibility, RS-FC of PrC and regions associated with the DMN and DAN is linked to worse performance in cognitive flexibility tasks. Taking our previous findings into account, a similar RS-FC map was positively associated with age, which could be because of a dedifferentiation in activation patterns as proposed in the dedifferentiation theory of neural aging (Baltes & Lindenberger, 1997; Goh, 2011; Li & Sikström, 2002; Park et al., 2001; 2004) or compensatory activations as postulated in CRUNCH (Reuter-Lorenz & Cappell, 2008), and STAC (Park & Reuter-Lorenz, 2008). However, given the nature of the available data and the methods applied, we cannot draw firmer and more theory-specific conclusions.

Summing up, our findings suggest that left aC/PrC is specifically recruited by older (vs. younger) adults, possibly to compensate for difficulties in shifting their attentional focus. Conversely, our results indicate an age-related increase in relative aC/PrC deactivation during the control task and/or an age-related decrease in relative aC/PrC deactivation during the experimental task, rising an alternative hypothesis for the higher task-induced aC/PrC activation in older adults. Left aC/PrC’s intrinsic coupling with the DMN and DAN supports its proposed role as a specialized hub, involved in internally as well as externally oriented information processing. The age-related decrease in RS-FC between aC/PrC and sensorimotor networks suggests some decoupling with age that is detrimental to action-related, externally oriented processing; the concurrent increase in RS-FC between DMN and DAN, in turn, suggests age-related difficulties in decoupling aC/PrC from the DMN during task states and from DAN-related regions during rest. Taking left aC/PrC’s often reported covariation with left IFJ during rest into account which was not found in the current study, our findings might reflect (and possibly contribute to) a dediffer-entiation in functional network patterns in older adults, potentially undermining the special role this region plays in shifting between internally and externally directed attention.

### 4.4 Limitations and Outlook

Although ALE is a well validated and widely used coordinate-based meta-analytic approach, it stands to reason that image-based meta-analyses may have provided greater sensitivity (Salimi-Khorshidi et al., 2009). However, as imaging data have previously been rarely shared, it would have been difficult to impossible to find a sufficient number of experiments with wholebrain images of effect estimates and standard errors.

Further, we were not able to conduct domain-specific meta-analyses for working memory and cognitive flexibility, since too few experiments were eligible for inclusion. More individual fMRI studies would be necessary to separately investigate the three EF subcomponents. The inclusion of more experiments would furthermore allow for testing a domain-specific account of EFs by directly contrasting the subcomponents with each other and testing additional or different EF subdivisions including even more fine-grained EF subprocesses. As previously discussed in the context of left IFJ and left aC/PrC, it seems that there is a process-specific sensitivity to aging. This process specificity may strongly contribute to the observed small to nonexistent across-experiment convergence of age differences in regional EF-related brain activity. In the context of inhibitory control, Korsch et al. (2014) found different age effects for different conflict tasks. In particular, there was overlap in brain activation during a flanker task between the two age groups and additional age-related activity in parietal and frontal regions. In contrast, during a stimulus– response compatibility task, no overlap in brain activation between the two groups was observed. Hence, age differences in EF-related brain activity appear to be task-specific to a substantial degree. This, in turn, would then lead to a heterogeneous distribution of age-related effects across studies, even within EF subdomains, which severely limits the chances for meta-analytic convergence and would argue for changing the focus of future research away from attempting to localize common, (sub)domain-general activation differences between age groups toward identifying process-specific mechanisms of age-related activity modulations. As discussed earlier, another explanation could be age-related regional changes in grey-matter volume (i.e. atrophy). Thus, we recommend that future studies on this topic investigate (i) domains and even subdomains, (ii) compare age-related differences in EFs across different modalities, and (iii) incorporate computational cognitive modeling (Kriegeskorte & Douglas, 2018).

Additionally, due to the small number of studies that reported deactivations, we were only able to investigate activation effects. As our results indicate age-related difficulties in deactivating left aC/PrC in the context of EF-tasks, we call for future studies investigating both directions of task-induced brain activity changes.

Somewhat surprisingly, no significant correlations between the two seeds’ whole-brain RS-FC patterns and the EF subcomponents working memory and inhibition were found. As there is ample evidence for RS-FC correlations with EF abilities in the literature (e.g., Hampson et al., 2006; Markett et al., 2013), a possible explanation could be that the tests used to assess EF domain-related abilities (via compound scores) were not sufficiently representative of the rather broad EF subdomains to yield a valid assessment of individual abilities or, the breadth of the subdomains prevented the scores from sufficiently reflecting particular subprocesses and age modulations thereof. The latter notion is supported by the fact that age correlated only moderately with the combined EF score (*r* = -.44, *p* < .001), the cognitive flexibility score (*r* = -.41, *p* < .001), and the inhibitory control score (r = -.31, p < .001). It did only weakly correlate with the working memory score (*r* = -.15, *p* < .05). For future studies on these questions it may be beneficial to incorporate various psychometric assessments of a particular cognitive function, which would allow isolating function- and test-specific variance in order to elucidate brain–behavior relationships (and their changes across the lifespan) at a more commensurate level of “granularity”.

Comparing our results to those of earlier neuroimaging meta-analyses of age-related differences in EFs underlines the importance of (i) transparently reporting the analysis choices made, (ii) providing a detailed description of inclusion and exclusion criteria and their motivation, and (iii) precisely reporting the papers and contrasts included as well as whether further information was received from the authors of the original study (for guidelines see Müller et al., 2018). Otherwise, even meta-analyses lack comparability and reproducibility.

### 4.5 Conclusion

The current study suggests that left IFJ and left aC/PrC play an important role in age-related differences in EFs as they were found the only two brain regions that showed consistent age differences in their recruitment during EF tasks across three major domains (working memory, inhibitory control, and cognitive flexibility). Although RS-FC analyses point towards a domain-general role of left IFJ in EFs, the pattern of contributions to the meta-analytic results also suggests process-specific modulations by age. In particular, older adults appear to rely more on left IFJ in the context of cognitive flexibility and inhibition, whereas younger adults recruited it more strongly in the context of working memory. Our findings further indicate that left aC/PrC is specifically recruited by older adults during EF tasks, potentially reflecting inefficiencies in switching the attentional focus. Overall, our results question earlier meta-analytic findings that suggested different and more comprehensive sets of brain regions as showing consistent age modulations of their EF-related activity. Rather, our findings attest to the substantial heterogeneity of such age-related differences and call for research that pays more attention to replicability and focuses on more narrowly and precisely defined EF subprocesses by combining multiple behavioral assessments, computational cognitive modelling, and multi-modal imaging.

## 5. Acknowledgements

We thank all contacted authors who contributed results of relevant contrasts not explicitly reported in the original publications, and we apologize to all authors whose eligible papers we might have missed.

## 9. Appendix

**Table A1.**
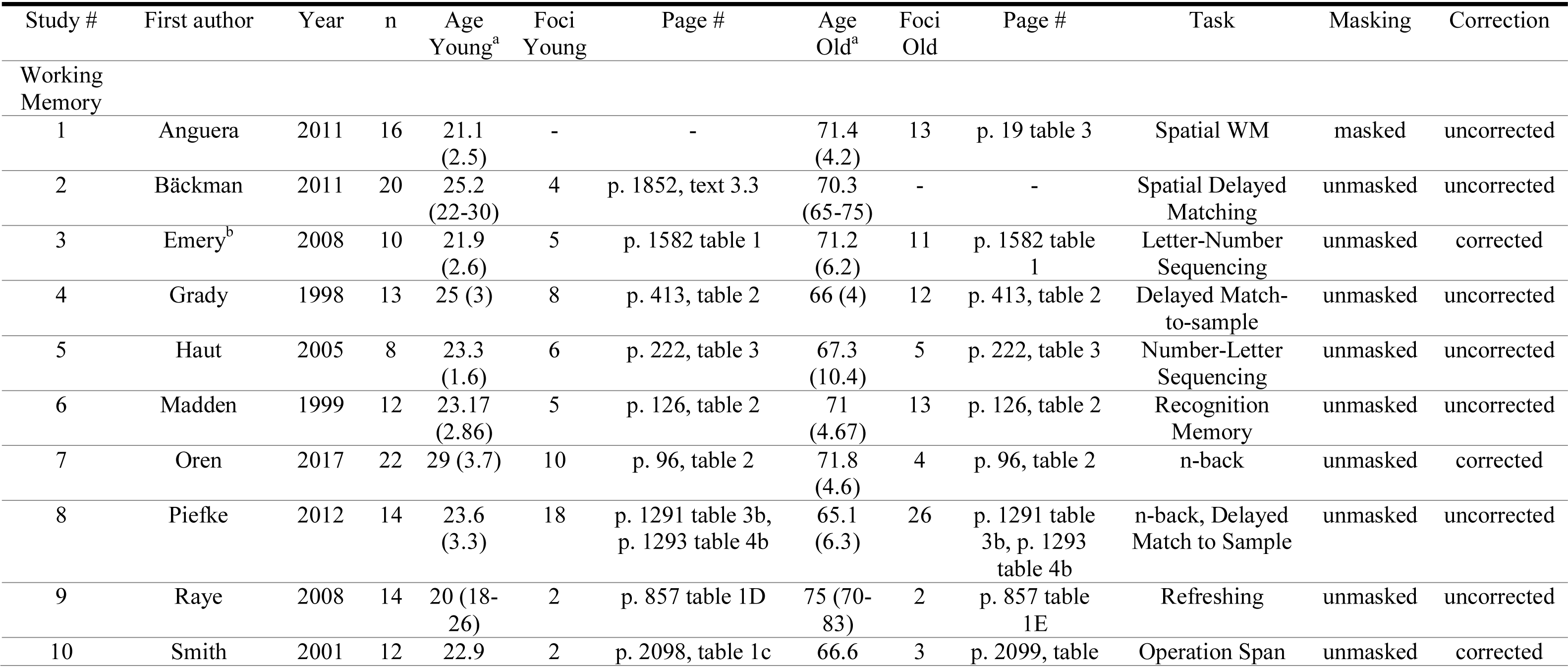
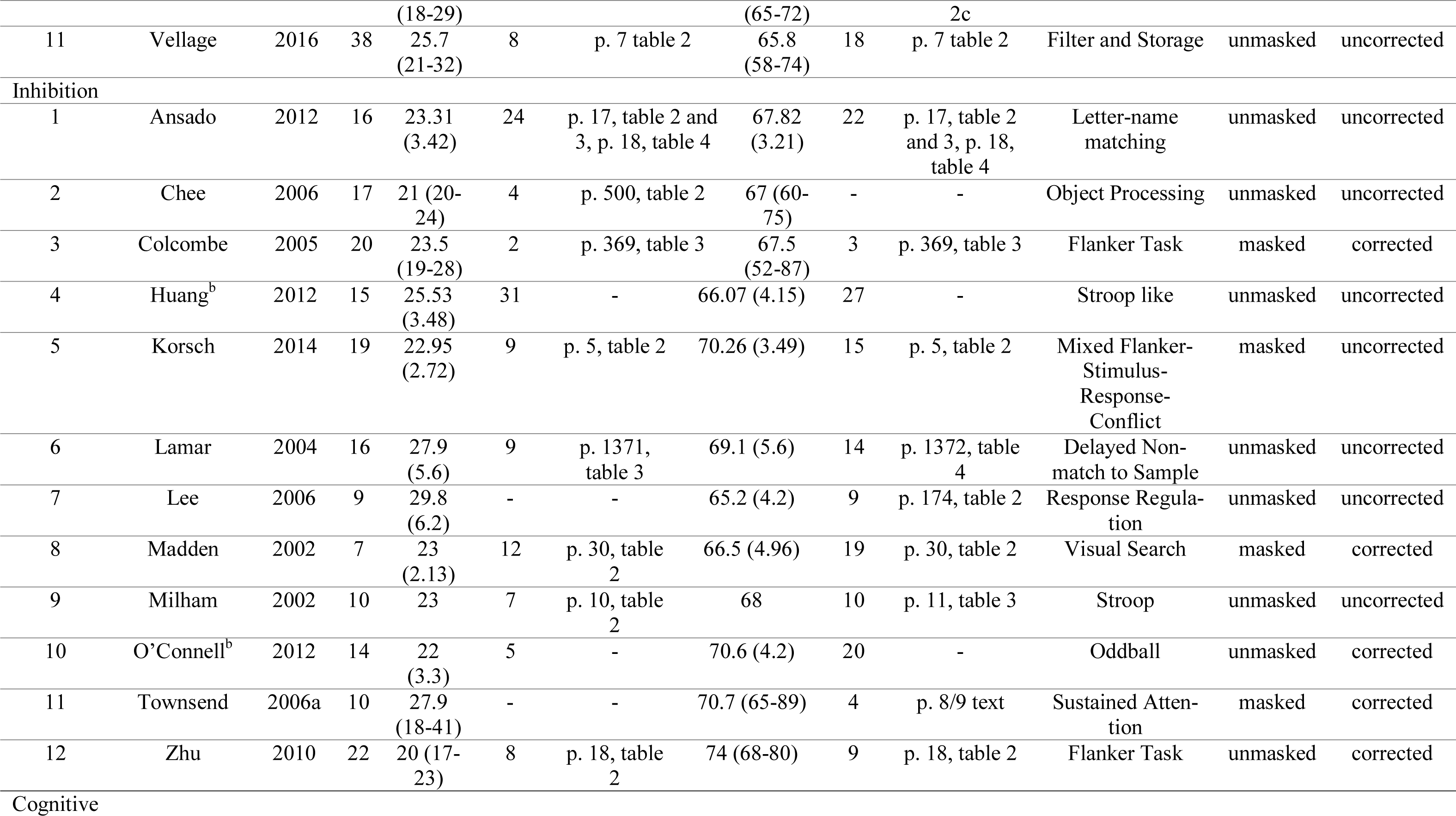
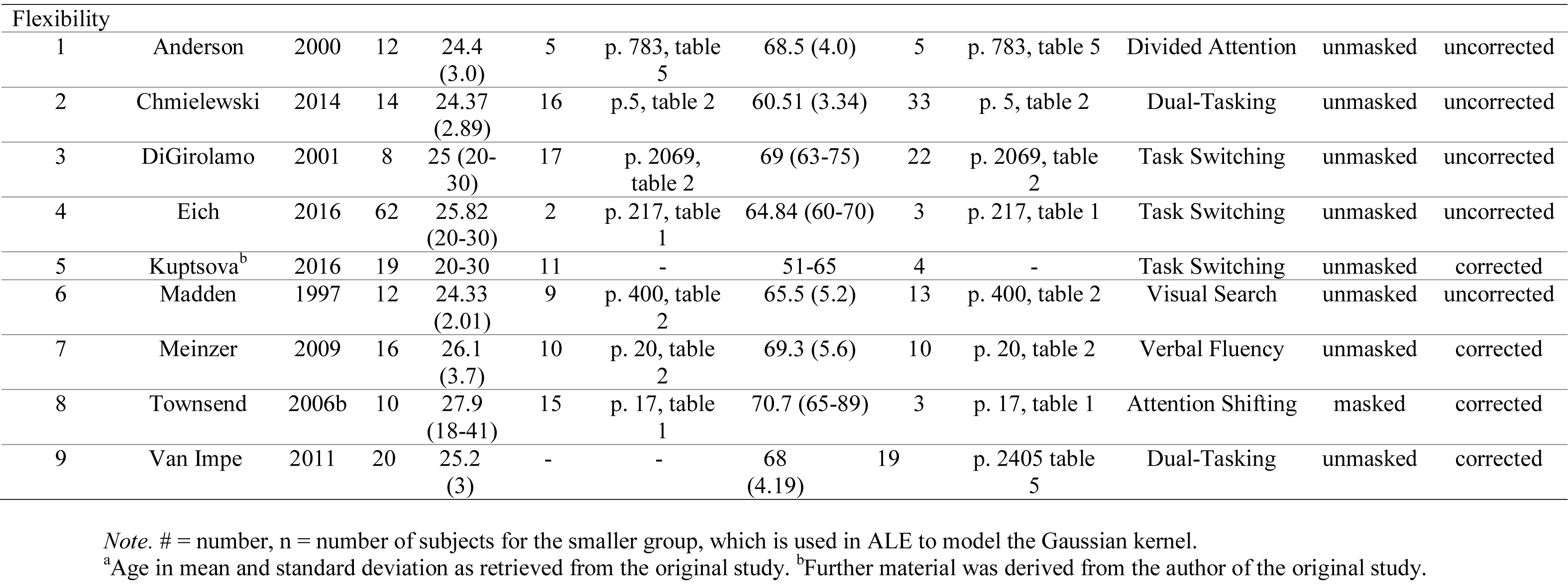
Overview of All Studies Included in the Meta-Analysis of Within-Group Contrasts Comprising Information About the Mean Age, Number of Activation Foci for Each Age Group, Masking with Task-Positive Effect, and Correction

**Table A2.** Overview of All Studies Included in the Meta-Analyses of Between-Group Contrasts Comprising Information About the Mean Age, Number of Activation Foci for Each Age Group, Masking with Task-Positive Effect, and Correction

| Study # | First author | Year | n | Age<br>Young<sup>a</sup> | Foci<br>Y>O | Page # | Age<br>Old<sup>a</sup> | Foci<br>O>Y | Page # | Task | Masking | Correction |
| --- | --- | --- | --- | --- | --- | --- | --- | --- | --- | --- | --- | --- |
| Working Memory |  |  |  |  |  |  |  |  |  |  |  |  |
| 1 | Anguera | 2011 | 16 | 21.1<br>(2.5) | 1 | p. 20 table 5 | 71.4<br>(4.2) | 10 | p. 20 table 5 | Spatial WM | masked | uncorrected |
| 2 | Bäckman | 2011 | 20 | 25.2<br>(22-30) | 3 | p. 1852 text 3.3 | 70.3<br>(65-75) | 4 | p. 1852 text 3.3 | Spatial Delayed Matching | unmasked | uncorrected |
| 3 | Bennett | 2013 | 20 | 21.8<br>(2.5) | 1 | supplementary table 3 | 65.3<br>(5.3) | 3 | supplementary table 2 | Delayed Item Recognition | masked | corrected |
| 4 | Emery<sup>b</sup> | 2008 | 10 | 21.9<br>(2.6) | - | - | 71.2<br>(6.2) | 37 | supplementary table 1 | Letter-Number Sequencing | unmasked | corrected |
| 5 | Fakhri | 2012 | 16 | 21 (3.7) | 5 | p. 358 table 3 | 68 (7.9) | 5 | p. 358 table 3 | Probe Recognition | unmasked | corrected |
| 6 | Grady | 1998 | 13 | 25 (3) | 4 | p. 413 table 2, p. 418 figure 4C, p. 419 table 4 | 66 (4) | 2 | p. 413 table 2, p. 418 figure 5C | Delayed Match-to-sample | unmasked | uncorrected |
| 7 | Grady | 2007 | 16 | 26.1<br>(3.7) | - | - | 65.8<br>(4.5) | 8 | p. 196 table 4 | N-back | unmasked | uncorrected |
| 8 | Haut | 2005 | 8 | 23.3<br>(1.6) | 1 | p. 222 table 4 | 67.3<br>(10.4) | - | - | Number-Letter Sequencing | unmasked | uncorrected |
| 9 | Kurth | 2016 | 20 | 23.4<br>(8.7) | 2 | p. 89 table 4 | 74.4<br>(5.6) | 10 | p. 89 table 4, p. 90 table 5 | Probe Recognition | unmasked | corrected |
| 10 | Lamar | 2004a | 16 | 27.9<br>(5.6) | 13 | p. 1372 table 5 | 69.1<br>(5.6) | 4 | p. 1372 table 5 | Delayed Match-to-sample | unmasked | uncorrected |
| 11 | LeCouvey | 2015 | 34 | 46.79<br>(18.82) | 3 | p. 7 table 4A | 46.79<br>(18.82) | - | - | Binding Task | masked | uncorrected |
| 12 | Paxton<sup>b</sup> | 2007a | 20 | 22.8<br>(3.7) | 9 | p. 34 table 2,<br>supplementary<br>table 2b | 73 (5.7) | 22 | p. 34 table 2,<br>supplementary<br>table 2b | AX-CPT | unmasked | uncorrected |
| 13 | Podell | 2012 | 11 | < 35 | 16 | supplementary<br>table 2 | > 65 | - | - | Updating WM | masked | corrected |
| 14 | Prakash | 2012 | 25 | 23.4<br>(3.3) | 4 | p. 195 table 2 | 72.16<br>(4.6) | - | - | N-back | masked | corrected |
| 15 | Raye | 2008 | 14 | 20 (18-26) | 2 | p. 857 table 1D | 75 (70-83) | 2 | p. 857 table<br>1E | Refreshing | unmasked | uncorrected |
| Inhibition |  |  |  |  |  |  |  |  |  |  |  |  |
| 1 | Ansado | 2012 | 16 | 23.31<br>(3.42) | 4 | p. 17 table 3 and<br>4 | 67.82<br>(3.21) | 9 | p. 17 table 2<br>and 3 | Letter-name<br>matching | unmasked | uncorrected |
| 2 | Bloemen-<br>daal | 2016 | 23 | 22.7<br>(0.6) | 2 | supplementary<br>figure 7 | 67.6<br>(0.7) | 9 | supplementary<br>table 5 | Load Depend-<br>ent Stop-signal<br>Anticipation | unmasked | uncorrected |
| 3 | Dørum | 2016 | 21 | 24.42<br>(5.06) | 6 | p. 7 table 2 | 64.67<br>(7.44) | - | - | Multiple Ob-<br>ject Tracking | unmasked | corrected |
| 4 | Eich | 2016a | 62 | 25.82<br>(20-30) | - | - | 64.84<br>(60-70) | 6 | p. 218 table 3,<br>p. 219 table 4 | Task switching<br>with go/no-go<br>component | unmasked | uncorrected |
| 5 | Grady | 2010 | 10 | 25 (3) | 10 | p. 173 table 3 | 66 (4) | 18 | p. 173 table 3 | Face Discrimi-<br>nation | unmasked | uncorrected |
| 6 | Huang | 2012 | 15 | 25.53<br>(3.48) | - | - | 66.07<br>(4.15) | 16 | p. 26 table 4 | Stroop like | unmasked | uncorrected |
| 7 | Korsch | 2014 | 19 | 22.95<br>(2.72) | 1 | p. 7 table 3 | 70.26<br>(3.49) | 3 | p. 7 table 3 | Mixed Flanker-Stimulus-Response-Conflict | masked | uncorrected |
| 8 | Lamar | 2004 | 16 | 27.9 | 6 | p. 1372 table | 69.1 | 8 | p. 1372 table | Delayed Non- | unmasked | uncorrected |

|  |  | b |  | (5.6) |  | 5 | (5.6) |  | 5 | match to Sam-<br>ple |  |  |
| --- | --- | --- | --- | --- | --- | --- | --- | --- | --- | --- | --- | --- |
| 9 | Lange-<br>necker | 2004 | 13 | 26.3<br>(5.5) | - | - | 71.1<br>(5.4) | 22 | p. 196 table 4<br>and 5 | Stroop | unmasked | corrected |
| 10 | Lee | 2006 | 9 | 29.8<br>(6.2) | - | - | 65.2<br>(4.2) | 3 | p. 174 table 2 | Response<br>Regulation | unmasked | uncorrected |
| 11 | Madden | 2002 | 7 | 23<br>(2.13) | 5 | p. 30 table 2 | 66.5<br>(4.96) | - | - | Visual Search | masked | corrected |
| 12 | Milham | 2002 | 10 | 23 | 6 | p. 10 table 2 | 68 | 4 | p. 11 table 3 | Stroop | unmasked | uncorrected |
| 13 | O'Connell | 2012 | 14 | 22 (3.3) | - | - | 70.6<br>(4.2) | 2 | p. 9 table 4 | Oddball | unmasked | corrected |
| 14 | Paxton<sup>b</sup> | 2007<br>b | 16 | 21.56<br>(3.14) | 1 | supplemen-<br>tary table 5b | 72.38<br>(6.51) | 29 | p. 36 table 3,<br>supplementary<br>table 5b | AX-CPT | unmasked | uncorrected |
| 15 | Persson<sup>b</sup> | 2007 | 28 | 21.7<br>(2.5) | - | - | 68.1<br>(5.8) | 4 | received from<br>author | Verb Genera-<br>tion | masked | corrected |
| 16 | Schulte | 2011 | 14 | 23.6<br>(19-30) | 9 | p. 2083 table<br>2, p. 2084<br>table 3 | 71 (58-85) | 16 | p. 2083 table<br>2, p. 2084<br>table 3 | Stroop Match-to-sample | unmasked | uncorrected |
| 17 | Sebastian | 2013 | 49 | 39.96<br>(17.14) | 8 | p. 2188 table<br>3 | 39.96<br>(17.14) | 12 | p. 2188 table<br>3 |  | masked | corrected |
| 18 | Townsend | 2006a | 10 | 27.9<br>(18-41) | - | - | 70.7<br>(65-89) | 4 | p. 9 text | Sustained At-<br>tention | masked | corrected |
| 19 | Zysset | 2007 | 23 | 26.6<br>(3.6) | - | - | 57.1<br>(6.49) | 7 | p. 941 table 2 | Stroop | unmasked | uncorrected |
| Cognitive<br>Flexibil-<br>ity |  |  |  |  |  |  |  |  |  |  |  |  |
| 1 | Chmielews<br>ki | 2014 | 14 | 24.37<br>(2.89) | - | - | 60.51<br>(3.34) | 2 | p. 193 text | Dual-Tasking | unmasked | uncorrected |
| 2 | Eich | 2016<br>b | 62 | 25.82<br>(20-30) | 7 | p. 217 table 1 | 64.84<br>(60-70) | 21 | p. 217 table 1 | Task Switching | unmasked | uncorrected |
| 3 | Fernandes | 2006 | 11 | 26.33<br>(3.36) | 2 | p. 2459 table<br>5 | 71.18<br>(4.07) | 8 | p. 2459 table<br>5 | Divided Atten-<br>tion | masked | uncorrected |
| 4 | Hubert | 2009 | 12 | 22.4<br>(2.5) | - | - | 65 (4.5) | 6 | p. 15 table 5 | Task of To-<br>ronto | masked | uncorrected |
| 5 | Kunimi | 2016 | 20 | 23.85<br>(5.43) | - | - | 67.35<br>(4.27) | 22 | p. 23 table 2 | Task Switching | unmasked | corrected |
| 6 | Kuptsova | 2016 | 19 | 20-30 | - | - | 51-65 | 29 | p. 367 table 3 | Task Switching | unmasked | corrected |
| 7 | Madden | 1997 | 12 | 24.33<br>(2.01) | 5 | p. 400 and<br>401 table 2 | 65.5<br>(5.2) | 5 | p. 400 and<br>401 table 2 | Visual Search | unmasked | uncorrected |
| 8 | Madden | 2010 | 20 | 22.4<br>(2.5) | 1 | p. 36 table 3 | 69.6<br>(6.05) | 17 | p. 36 table 3 | Task Switching | masked | uncorrected |
| 9 | Meinzer | 2009 | 16 | 26.1<br>(3.7) | - | - | 69.3<br>(5.6) | 5 | p. 8 text | Verbal Fluency | unmasked | corrected |
| 10 | Steffener<sup>b</sup> | 2016 | 63 | 25.79<br>(2.7) | - | - | 65.47<br>(2.89) | 13 | received from<br>author | Task Switching | masked | uncorrected |
| 11 | Townsend | 2006<br>b | 10 | 27.9<br>(18-41) | 3 | p. 18 table 2 | 70.7<br>(65-89) | 20 | p. 18 table 2 | Attention<br>Shifting | masked | corrected |
| 12 | Van Impe | 201<br>1 | 20 | 25.2 (3) | - | - | 68<br>(4.19) | 19 | p. 2405 table<br>5 | Dual-Tasking | unmasked | corrected |
| 13 | Zhu | 201<br>6 | 28 | 32 (3.8) | - | - | 68.4<br>(5.4) | 18 | p. 141 table 3 | Task Switching | unmasked | corrected |
| 14 | Worthy | 201<br>6 | 18 | 23.61<br>(18-31) | 6 | p. 18 table 2 | 67 (61-79) | - | - | Decision Mak-<br>ing | unmasked | corrected |
| Miscalle-<br>neous |  |  |  |  |  |  |  |  |  |  |  |  |
| 1 | Esposito | 199<br>9 | 41 | 45.5<br>(19.7) | 13 | p. 969 table 1,<br>p.970 table 2 | 45.5<br>(19.7) | 11 | p. 969 table 1,<br>p.970 table 2 | WCST, RPM | unmasked | uncorrected |

*Note.* # = number, n = number of subjects for the smaller group, which is used in ALE to model the uncertainty of coordinates, Y>O = young > old, O>Y = old > young.
<sup>a</sup>Age in mean and standard deviation as retrieved from the original study. <sup>b</sup>Further material was obtained from the authors of the original study.

**Table A3.** Checklist for Neuroimaging Meta-Analyses by Müller et al. (2018)

|  |  |
| --- | --- |
| The research question was specifically defined | YES, and it included the following contrasts:<br>1) Within-group contrasts for young<br>2) Within-group contrasts for old<br>3) Between-group contrasts young > old<br>4) Between-group contrasts for old > young<br>The specific contrasts are reports in Tables 1 and 2 |
| The literature search was systematic | YES, it included the following keywords in the following databases:<br>1) (1) title: “age” or “aging” or “ageing” or “age-related” or “older adults” or “old adults” or “life-span” or “elderly adults”; and (2) title: “executive functions” or “working memory” or “inhibition” or “cognitive flexibility”; and (3) abstract: “fMRI” or “functional magnetic resonance imaging” or “PET” or “positron emission tomography” or “neuroimaging” or “cerebral blood flow”<br>2) For working memory “n-back” or “sternberg” or “delayed match to sample” or “delayed simple matching” or “stimulus-response-compatibility”; for inhibition “stroop” or “flanker” or “simon” or “stop signal” or “go/no-go” or “stimulus detection” or “stimulus discrimination” or “selective attention”; for cognitive flexibility “task switching” or “dual task” or “set shifting”<br>3) Databases: Web of Science (http://apps.webofknowledge.com), PubMed (https://www.ncbi.nlm.nih.gov/pubmed/), PsycINFO (http://ovidsp.tx.ovid.com), and Google Scholar (http://scholar.google.de) |
| Detailed inclusion and exclusion criteria were applied | YES, and reasons of non-standard criteria were:<br>Inclusion of: |
|  | - fMRI and PET studies<br>- Healthy young and old participants without any pharmacological manipulations<br>- Masking of the between-group contrast with task-positive effect<br>- Activation data<br>- For meta-analysis of within-group contrast: main task effect per group<br>- For meta-analysis of between-group contrast: group comparison, positive correlation with age (old > young), negative correlation with age (young > old)<br>- No correlation or interaction with other variables (e.g., RT)<br>- Task > sensorimotor control, task > resting-baseline, task difficult > task easy<br>- The difficult task condition was included, if contrasts representing easy and difficult conditions were available<br>- The contrast reflecting transient brain activity was included when contrasts for sustained and transient activity were available |
| Sample overlap was taken into account | YES, using the following method:<br>- Contribution from a study was limited to one experiment per study<br>- If a study reported several experiments eligible for inclusion, the reported coordinates were pooled to constitute a single experiment |
| All experiments used the same search coverage (state how brain coverage was assessed and how small volume corrections and conjunctions were taken into account) | YES, the search coverage was the following:<br>- Only whole-brain coverage<br>- Exclusion of ROI studies<br>- Inclusion of masking of the between-group contrast with task-positive effect |
| Studies are converted to a common reference space | YES, using the following conversion(s):<br>- Coordinates reported in Talairach space were converted to MNI space (Lancaster et al., 2007). |
| Data extraction was conducted by two investigators (ide- | YES, the following authors: |
| al case) or double-checked by the same investigator (state how double-checking was performed) | - Marisa Heckner, Edna Cieslik, and Robert Langner checked inclusion criteria<br>- Marisa Heckner extracted coordinates<br>- Marisa Heckner extracted other info: Number and age of subjects included, task, contrast, space, modality, masking of between-group contrast with task-positive effect, level of performance between age groups, correction of results<br>- Edna Cieslik double-checked the following data: Coordinates extracted, number and age of subjects included, task, contrast, space, modality, masking of between-group contrast with task-positive effect, level of performance between age groups, correction of results |
| The paper includes a table with at least the references, basic study description (e.g., for fMRI tasks, stimuli), contrasts and basic sample descriptions (e.g., size, mean age and gender distribution, specific characteristics) of the included studies, source of information (e.g., contact with authors), reference space | YES, and also the following data:<br>- If further information was received by the authors<br>- How coordinates were treated (MNI or Talairach) when space was not clearly specified in original study<br>- If the between-group contrast was masked with the task-positive effect<br>- If results were corrected for multiple comparisons<br>- Level of performance between age groups |
| The study protocol and all analyses was planned beforehand, including the methods and parameters used for inference, correction for multiple testing, etc. | YES:<br>1) No non-planned or post-hoc analyses.<br>2) The meta-analysis used the default methods and parameters of our group. |
| The paper includes meta-analytic diagnostics | Contributions from individual experiments to each cluster of significant convergence were provided for each meta-analysis performed. |

**Table A4.**
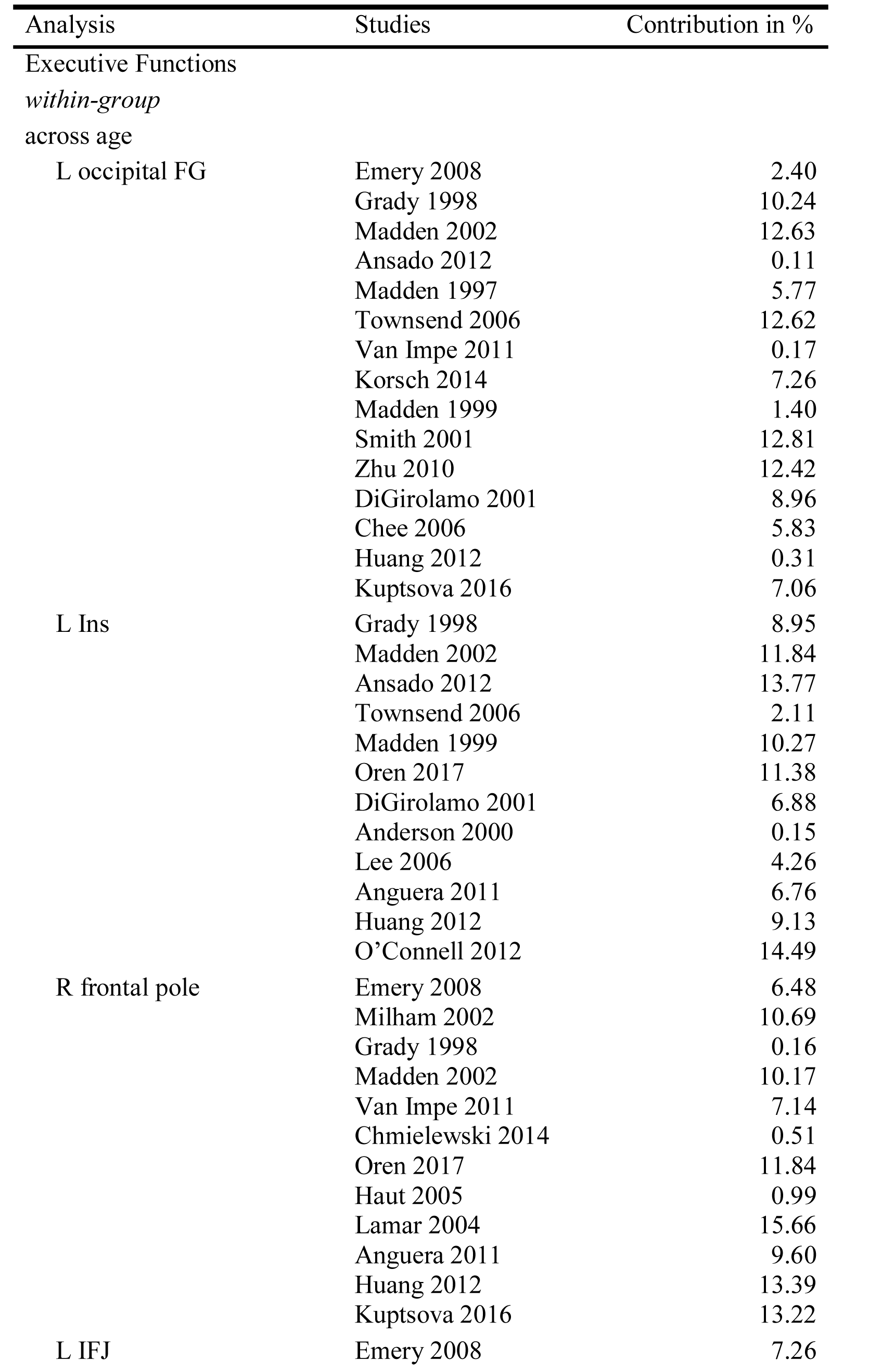
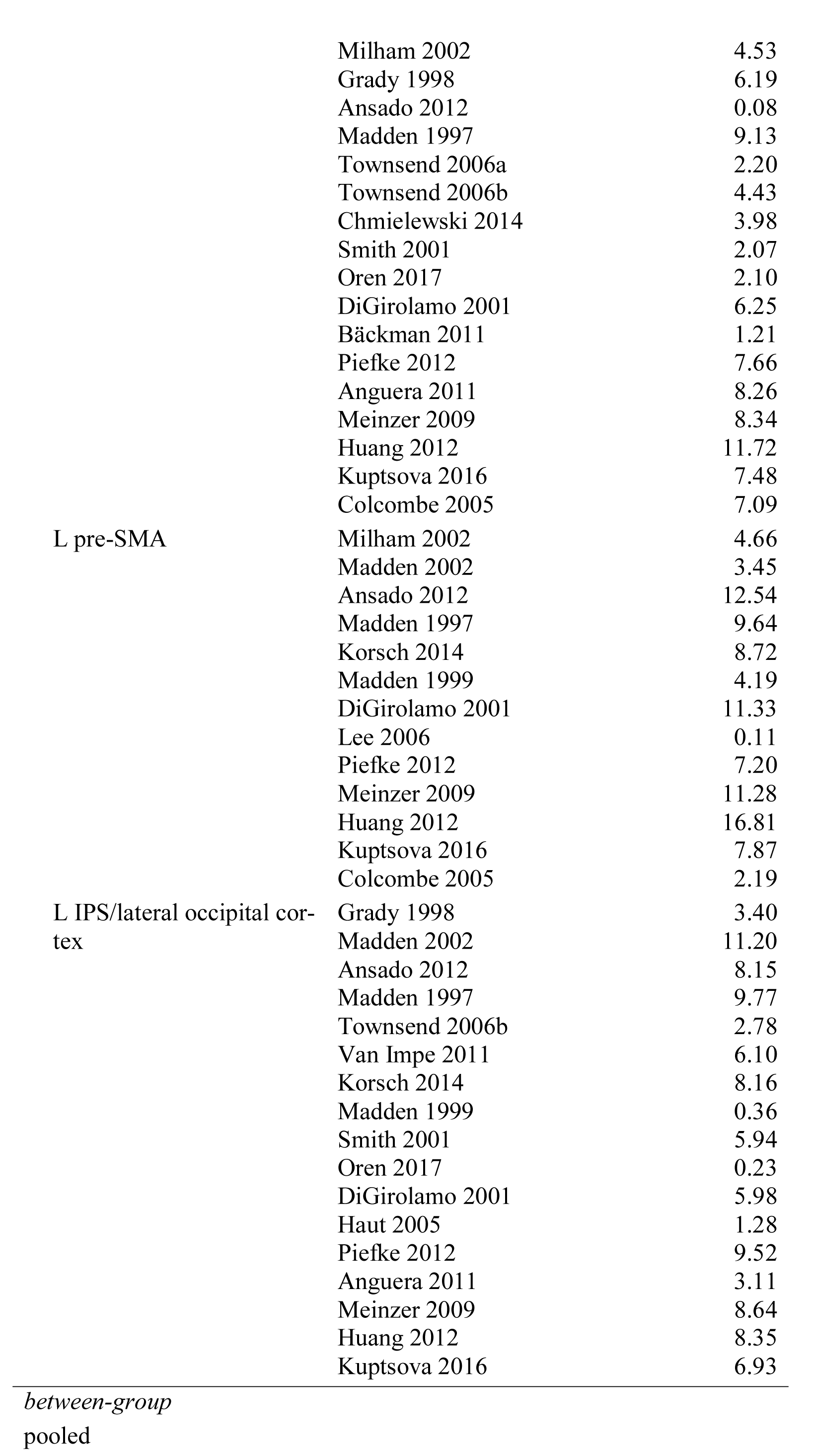
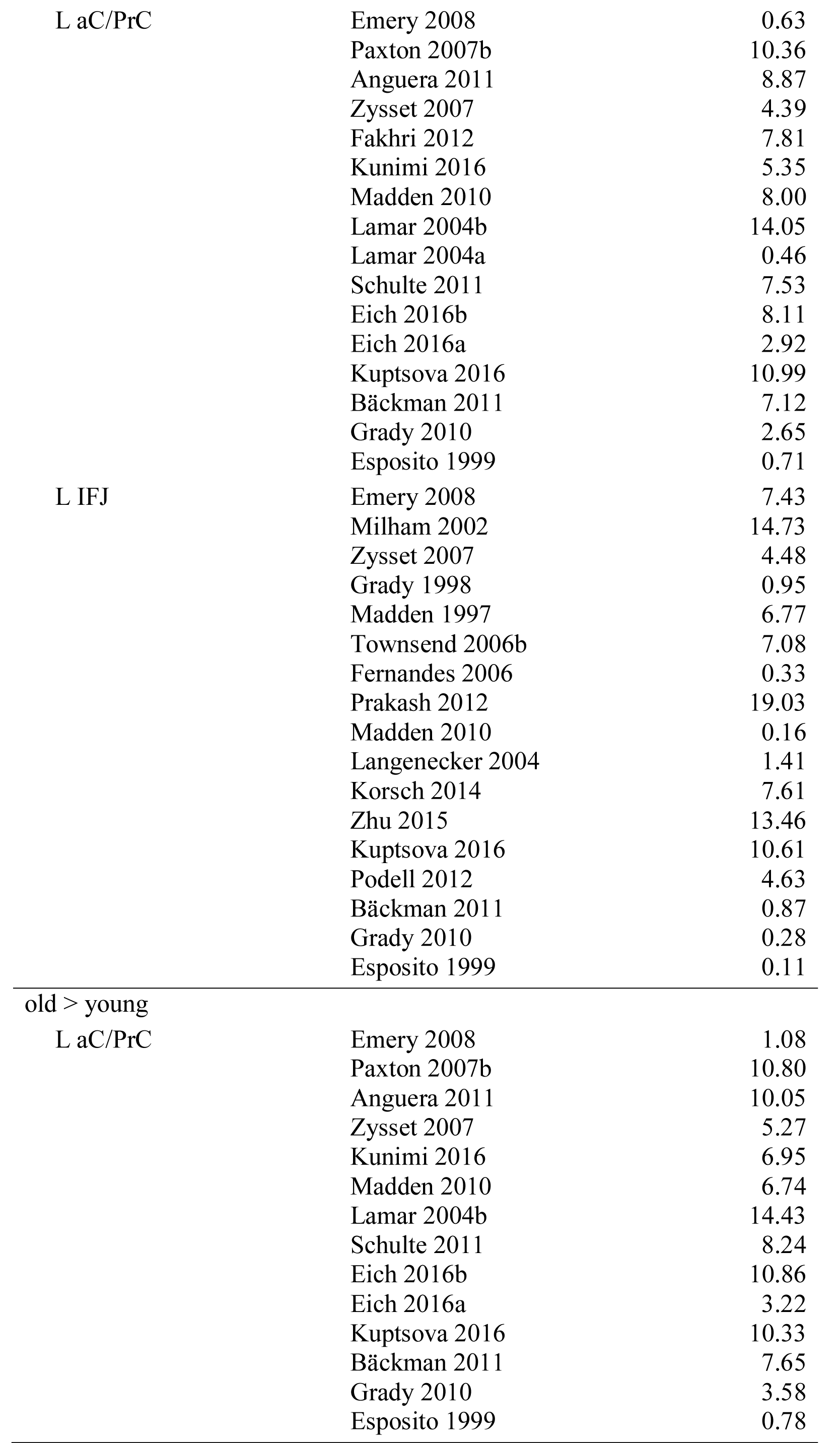
Single Experiments Contributing to the Clusters of Convergence

**Table A5.**
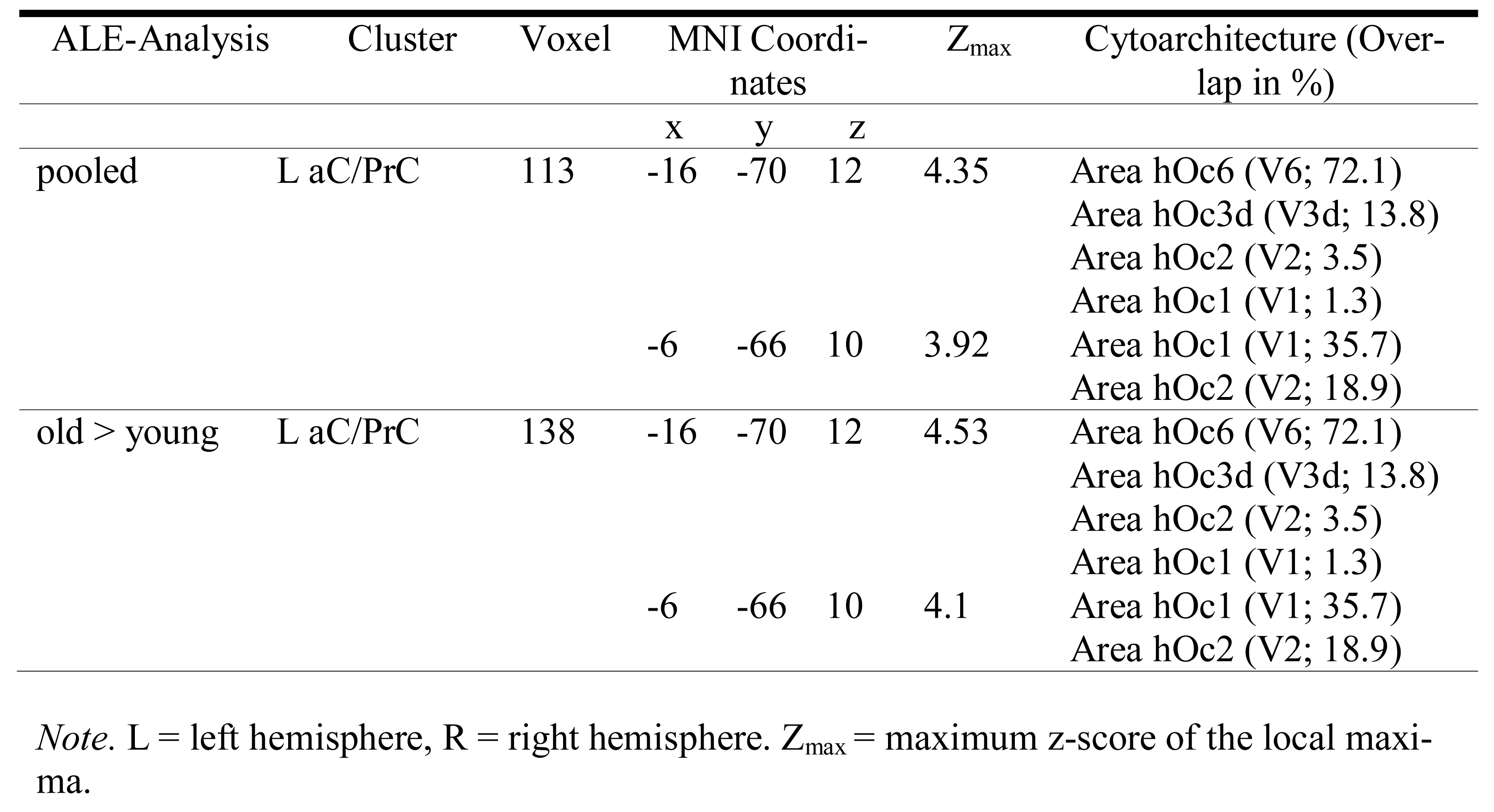
Brain Regions Showing Significant Convergence of Activity in Inhibition

**Table A6.** Single Experiments Contributing to the Inhibition Cluster of Convergence

| Analysis | Studies | Contribution in % |
| --- | --- | --- |
| pooled |  |  |
| L aC/PrC | Paxton 2007b | 20.97 |
|  | Zysset 2007 | 18.65 |
|  | Lamar 2004b | 30.08 |
|  | Schulte 2011 | 14.39 |
|  | Eich 2016a | 15.90 |
| old > young |  |  |
| L aC/PrC | Paxton 2007b | 21.83 |
|  | Zysset 2007 | 17.01 |
|  | Lamar 2004b | 29.46 |
|  | Schulte 2011 | 16.32 |
|  | Eich 2016a | 15.37 |

